# IndeCut evaluates performance of network motif discovery algorithms

**DOI:** 10.1101/156836

**Authors:** Mitra Ansariola, Molly Megraw, David Koslicki

**Author notes:** To whom correspondence should be addressed. Please address correspondence regarding sampling algorithms and their assessment in this work to Molly Megraw.: Please address correspondence regarding mathematics contained in this work to David Koslicki.

## Abstract

Genomic networks represent a complex map of molecular interactions which are descriptive of the biological processes occurring in living cells. Identifying the small over-represented circuitry patterns in these networks helps generate hypotheses about the functional basis of such complex processes. Network motif discovery is a systematic way of achieving this goal. However, a reliable network motif discovery outcome requires generating random background networks which are the result of a uniform and independent graph sampling method. To date, there has been no sound practical method to numerically evaluate whether any network motif discovery algorithm performs as intended—thus it was not possible to assess the validity of resulting network motifs. In this work, we present `IndeCut`, the first and only method that allows characterization of network motif finding algorithm performance on any network of interest. We demonstrate that it is critical to use `IndeCut` prior to running any network motif finder for two reasons. First, `IndeCut` estimates the minimally required number of samples that each network motif discovery tool needs in order to produce an outcome that is both reproducible and accurate. Second, `IndeCut` allows users to choose the most accurate network motif discovery tool for their network of interest among many available options. `IndeCut` is an open source software package and is available at <u>https://github.com/megrawlab/IndeCut</u>.

## 1 Introduction

Genomic networks represent a complex map of molecular interactions which are descriptive of the biological processes occurring in living cells [1, 2]. Due to the size and complexity of these networks, it is often difficult to infer the physiological function of individual interactions or collections of interactions without detailed additional information about network structure. Because this type of experimentally supported prior information is usually sparse or unavailable, a systematic approach for identifying key sub-components and their functions within a biological system is essential for analysis. From this perspective, it has been shown that the functional essence of a complex genetic network within a cell can often be distilled by thinking of the network as a “circuit board” composed of small, understandable components that work together to carry out higher-order processes [3, 4, 5, 6, 7, 2, 8, 9, 10]. Network motif discovery is a well-established statistical strategy for performing network analysis from this viewpoint. This strategy compares the frequency of observation of a sub-network within the larger original network to its frequencies in many randomized background networks in order to identify network motifs, which are defined as those sub-networks observed at a significantly higher frequency in the original network. In other words, a network motif is an over-represented sub-structure within a larger network.

Network motif discovery tools aid in generating specific testable hypotheses about the behavior and function of a genetic sub-circuit. For example, in the case of a gene regulatory network, a bi-stable switch coupled with a noise-damping circuit may be necessary to tune the expression of developmental transcription factors involved in body-plan patterning at a specific stage of development [3, 4, 11]; thus this circuit may appear as a motif in networks constructed from tissue samples in developing organisms. Although such hypotheses are valuable starting points for understanding the underlying mechanisms of a biological process through analysis of genomic networks, the laboratory validation of a predicted network motif is generally a costly and time-consuming endeavor. For example, validating a candidate regulatory sub-network containing a specific transcription factor, a microRNA, and a protein coding gene would typically require a series of procedures such as electrophoresis mobility shift assays and generation of reporter constructs, involving months of labor and thousands of dollars in supplies. This highlights the need for accurate network motif discovery procedures in order to acquire a biologically meaningful outcome.

Intuitively, identification of a biologically meaningful network motif results from a background network generation strategy that satisfies two conditions. 1) Background networks should preserve a sensible set of biological assumptions given the original network’s structure. For example, if the original network contains a node type (e.g. transcription factor) that can target itself as well as other genes in the original network, then the researcher may feel that this property should be preserved in the background networks. 2) Given these preserved properties, the background networks generated should provide a truly representative sample of all possible such networks. That is, the generation method should not favor the production of certain types of networks over others. While there are a variety of choices that a researcher may make about network property preservation, it is clearly desirable to generate an unbiased sample of background networks which preserve these properties-thus avoiding any hidden assumptions encoded in the background network generation procedure itself.

Computationally, the core component of background network generation is the process of sampling a number of networks (for example, 1000 networks) from the set of all possible networks (e.g. 1 million networks) having in-degree and out-degree sequences identical to those of the original biological network. In network motif algorithms, networks are usually thought of as graphs, and this sampling process is known as “graph sampling”. Ideally, graph sampling would be unnecessary; one would simply generate all possible graphs in the sample space, compute the number of a particular sub-graph of interest observed in each one, and then compute an exact P-value for this sub-graph by comparing the number of times it was observed in the original network to the probability of this number using the empirically identified background distribution. A very small P-value would indicate significant over-representation, and thus a network motif. Unfortunately, for networks of realistic biological size - even a few hundred nodes and edges - the size of the sample space is enormous (trillions of graphs). But more importantly, for the background network generation core component defined above there is no known closed-form formula for computing the number of such graphs. Thus, graph sampling is a practical necessity but presents a challenge in its own right, as one must sample in an unbiased manner from a set of unknown size.

Despite a rich mathematical literature on the subject [12, 13, 14, 15, 16, 17, 18, 19, 20], practical solutions to this problem remain elusive. As a result, to date, network motif discovery tools must perform graph sampling for sample spaces such that not only is uniformity and independence a concern, but the number of samples required to evaluate success/failure in conforming to this ideal is unknown. Several network motif discovery tools with different underlying graph sampling strategies are currently available [21, 22, 23]. Through experiments on smal“test graphs” where samples spaces can be em-pirically enumerated by producing all possible graphs in the space, it has been shown that the same sampling strategy can have very different performance outcomes in terms of uniform and independent sampling [3] depending on graph topology. For example, while d-regular graphs rarely pose a problem, small graphs with highly irregular or “uneven” degree sequences frequently cause difficulty [3, 24, 25]. This creates a severe concern for the accurate performance of network motif discovery algorithms on real biological networks, which often contain large source hubs (“master regulators”) and/or target hubs (heavily regulated nodes) [26, 27].

To date, no mathematically sound yet computationally practical method is available in order to determine whether a graph sampling method samples uniformly and independently for a large or even moderately-sized network of interest. Intuitively, estimating the degree of uniformity/independence of a set of sample graphs has remained a fundamental open challenge when the number of graphs in the complete sample space cannot be empirically determined. However, relatively recent advances in the enumerative combinatorics literature [28, 29] have opened an avenue for the development of solutions to this long-standing problem. In this study we present IndeCut, the very first practical method that determines the degree of sampling uniformity/independence for network motif discovery algorithms. IndeCut quantifies the performance of such algorithms by using a novel cut-norm based approach. IndeCut directly aids in understanding the cause of performance variations among different graph sampling approaches on a variety of biologically common graph topologies.

## 2 RESULTS

### 2.1 How does IndeCut work?

An ideal graph sampling strategy would produce samples from the set of all possible graphs (the *sample space*) that are perfectly uniform and independent. In the case of perfect sampling, each sample would have a sample average that is identical to the true average (mean of all elements in the sample space). From this perspective, violation of uniformity and independence can be quantified by measuring how far the *sample average* is from the *true average*. Figure 1 provides an abstracted visualization of this concept. Sampled graphs, described mathematically as zero-one matrices, are represented as gray dots inside the sample space of all graphs with the prescribed in and out-degrees. The point *E* represents the centroid, or true average, of the entire sample space and the point *A* represents the average of sampled matrices for a hypothetical graph sampling strategy. Figure 1A shows that a uniform and independent sampling method produces samples that are “evenly spread” over the sample space, resulting in a centroid matrix *E* and an average matrix *A* which are nearly identical. In the case of perfectly uniform and independent sampling, *E* and *A* are identical. In Figure 1, a violation of independence (Figure 1B) or uniformity (Figure 1C) results in a difference between *E* and *A*. The greater the violation of uniformity and/or independence, the further the centroid *E* will be from the sample average *A*.

**Figure 1:**
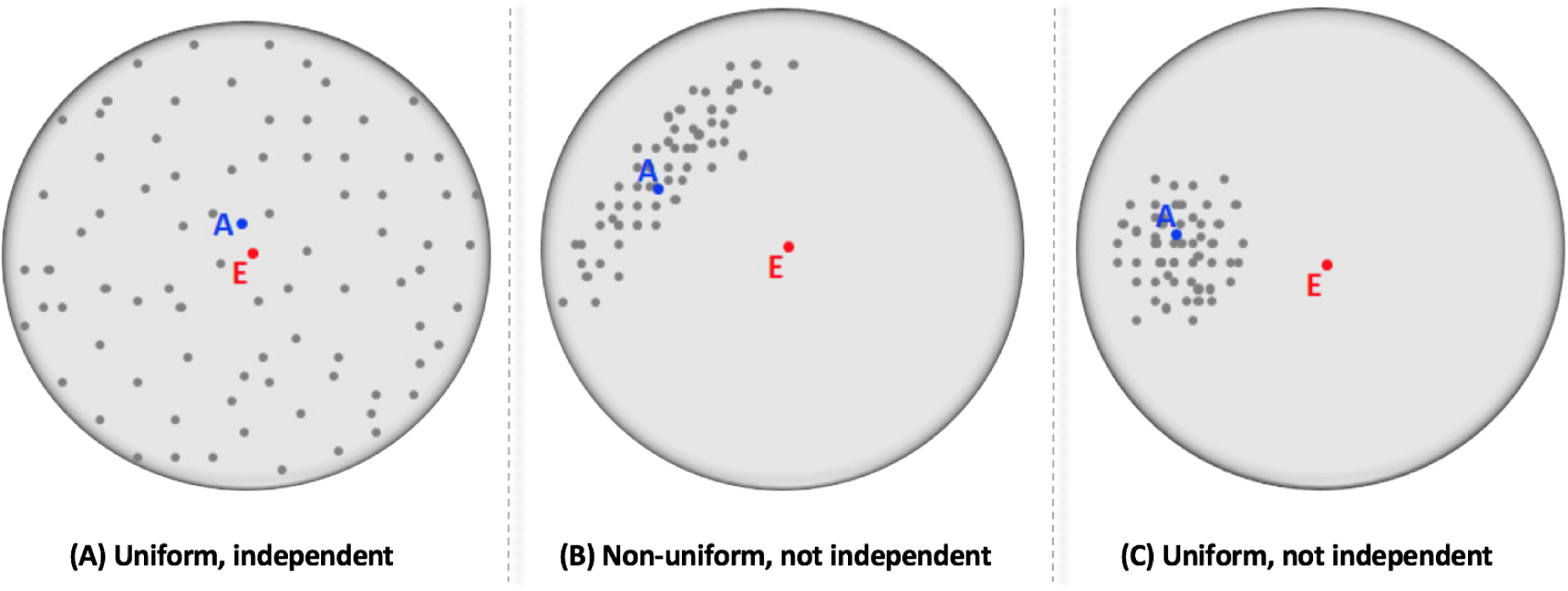
An illustrative view of graph sampling strategy outcomes in terms of uniformity and independence.

Computing the exact centroid *E* by empirically enumerating all graphs in the sample space is generally prohibitive because such spaces are astronomically large (for example, the space of 3-regular bipartite graphs with 10 source nodes and 10 target nodes has more than 10^26^ elements in it). Therefore instead of computing the exact centroid *E*, we use a proxy to this centroid (which we call the maximum entropy matrix and denote by *Z*) which is known to be “close” to *A* when the sampling regime is uniform and independent (i.e. a central limit theorem is satisfied for *Z* and proved by Barvinok [29]). The correct notion of “closeness” in this situation is a metric called the cut norm. Thus, an ideal sampling method will have a zero cut norm for *Z* - *A*, the matrix representing the difference between *Z* and *A*. Similarly, a cut norm bounded significantly away from zero indicates that sampling is either highly non-uniform, highly non-independent, or both. Unfortunately, computing the cut norm for matrices of realistic size is intractable given today’s computing hardware capability (MAX SNP-hard). We overcome this barrier by using the ideas of Alon and Naor [28] to create an approximation algorithm that returns an interval in which the distance between *Z* and *A* is guaranteed to be contained. Comparing these intervals allows us to compare the uniformity and independence of graph sampling strategies.

Here, we present IndeCut as a practical method that determines the degree of uniformity/independence for a sampling method on a given bipartite graph *G* with fixed in-degrees and out-degrees (and no multiple edges). *G* is a simple graph wherein source nodes have direct links to the target nodes but not vice-versa. The sample space associated with graph *G* is defined as the set of all possible valid bipartite graphs that can be produced from *G*’s degree sequence. Each bipartite graph in the sample space can be represented as a zero-one matrix with fixed row and column sums *R* and *C*, respectively (see Supplementary Figure S1).

In summary, IndeCut, performs the following tasks in a computationally efficient manner: the sample average matrix *A* and maximum entropy matrix *Z* are computed, and then a (typically small) interval is computed along with a guarantee that the cut norm lies in this interval. The further this interval is bounded away from zero, the less uniform and independent the graph sampling technique is. Importantly, no additional information is required by IndeCut to assess a whether a graph sampling strategy performs uniform/independent sampling other than the input graph and sequence of graphs returned by the sampling method. Further details, along with mathematically rigorous definitions and proofs, are provided in the Materials and Methods section.

### 2.2 IndeCut evaluates the performance of network motif discovery algorithms

IndeCut helps users to determine whether a network motif discovery algorithm is able to provide a valid network motif outcome (i.e. does it use a uniform/independent background network generation method) on a network of interest, for any algorithm of choice and any network topology. Prior to IndeCut, there was no method by which a user could determine which network motif discovery algorithm (if any) would provide an accurate/unbiased outcome on an input network. This posed a significant problem for biological networks in particular, given that experimental evaluation of such an outcome is costly and time-consuming. Here, we demonstrate how IndeCut can be used by examining several major graph topologies and commonly used algorithms. Two different types of graphs are examined: 1) small graphs with topologies that typically occur in biological networks, and 2) realistic graphs from the literature with a large number of nodes and edges. We selected four network motif discovery approaches from the recent literature: FANMOD (Fast Network Motif Detection) ([30]), DIA-MCIS (Diaconis Monte Carlo Importance Sampling) ([31]), WaRSwap (Weighted and Reverse Swap sampling) ([3]), and CoMoFinder (Coregulatory network Motif Finder) ([32]). Each of these algorithms represents a fundamentally different strategy for network motif discovery background network generation. The Methods Section provides a detailed description of each algorithm.

#### 2.2.1 Small graph collection

Three classes of small graphs were created to consider three distinct topological properties: 1) “uneven” (irregular) graphs containing large hub nodes (a hub node has a large in-degree or out-degree as compared to the other nodes in the graph). 2) “even” (regular) graphs with nearly even or even (d-regular) degree sequences. 3) “hybrid” combinations of in-degree and out-degree sequences that form graphs with various topological features (not limited to extreme cases of evenness or unevenness as in 1 and 2). These graphs mimic the properties of large biological networks on a smaller scale, and enable us to examine how IndeCut evaluates the sampling performance of different network motif discovery algorithms on specific graph structures. Supplementary Figure S1 shows the degree sequence of each graph examined.

For the uneven class, we created six hub-containing graphs. The first graph (uniFanG1) is an example of a simple hub-containing bipartite graph in which each layer (source and target layers) has a node with a large degree (out-degree and in-degree) as compared to other nodes in that layer. We increased the degree of unevenness of the uniFanG1 graph by joining another uni-fan graph to it to create a new graph with four large hubs (biFanG1). We repeated this process (attaching a uniFanG1 to an existing graph) to generate the “Fan” series of graphs (see Supplementary Figure S2). These graphs allow us to understand how IndeCut captures the performance of each algorithm on graphs with an increasingly large degree of unevenness, a topology type which is known to pose difficulties to many algorithms ([3]). For the class of regular graphs, three d-regular and three near d-regular graphs with a different number of nodes and edges were created and examined using IndeCut. Fig 2, Fig 3, and Fig 4 show the cut norm estimates for each graph and algorithm within three classes of small graphs. In each of these figures, 5000 random graphs were generated for each of the four methods, then cut norm bounds were computed using IndeCut. A smaller cut norm interval that is closer to zero represents more uniform and independent sampling.

**Figure 2:**
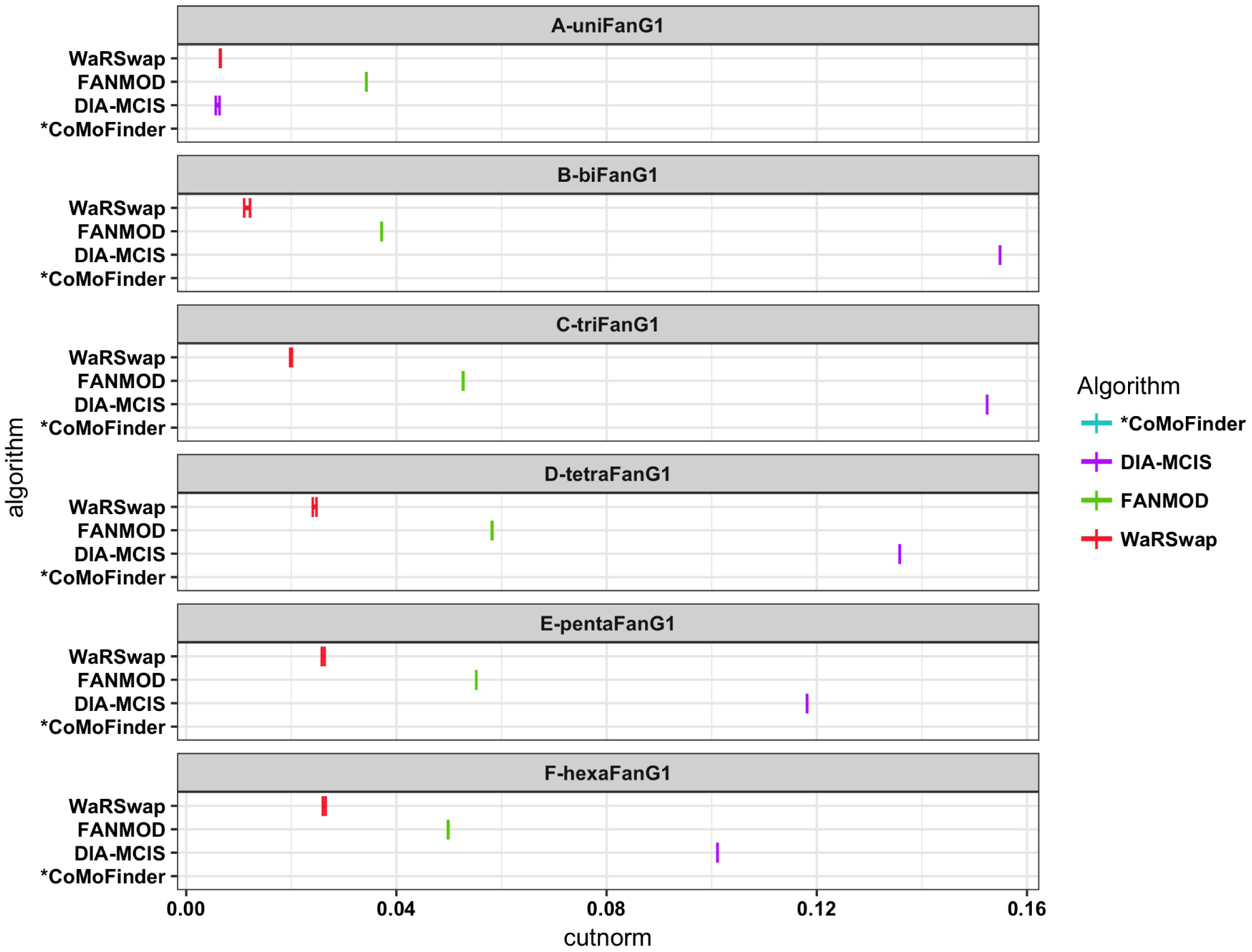
**Uniform/independent graph sampling performance evaluation on small uneven graphs,** for each small uneven graph and algorithm, 5000 graphs were generated. The cut norm estimates for each algorithm were computed using IndeCut. With the exception of uniFanG1, the cut norm estimates for CoMoFinder were much larger than 0.16, therefore we removed CoMoFinder’s results from this figure for ease of comparison (see Supplementary Table S1 and Supplementary Figure S3 for detailed results).

**Figure 3:**
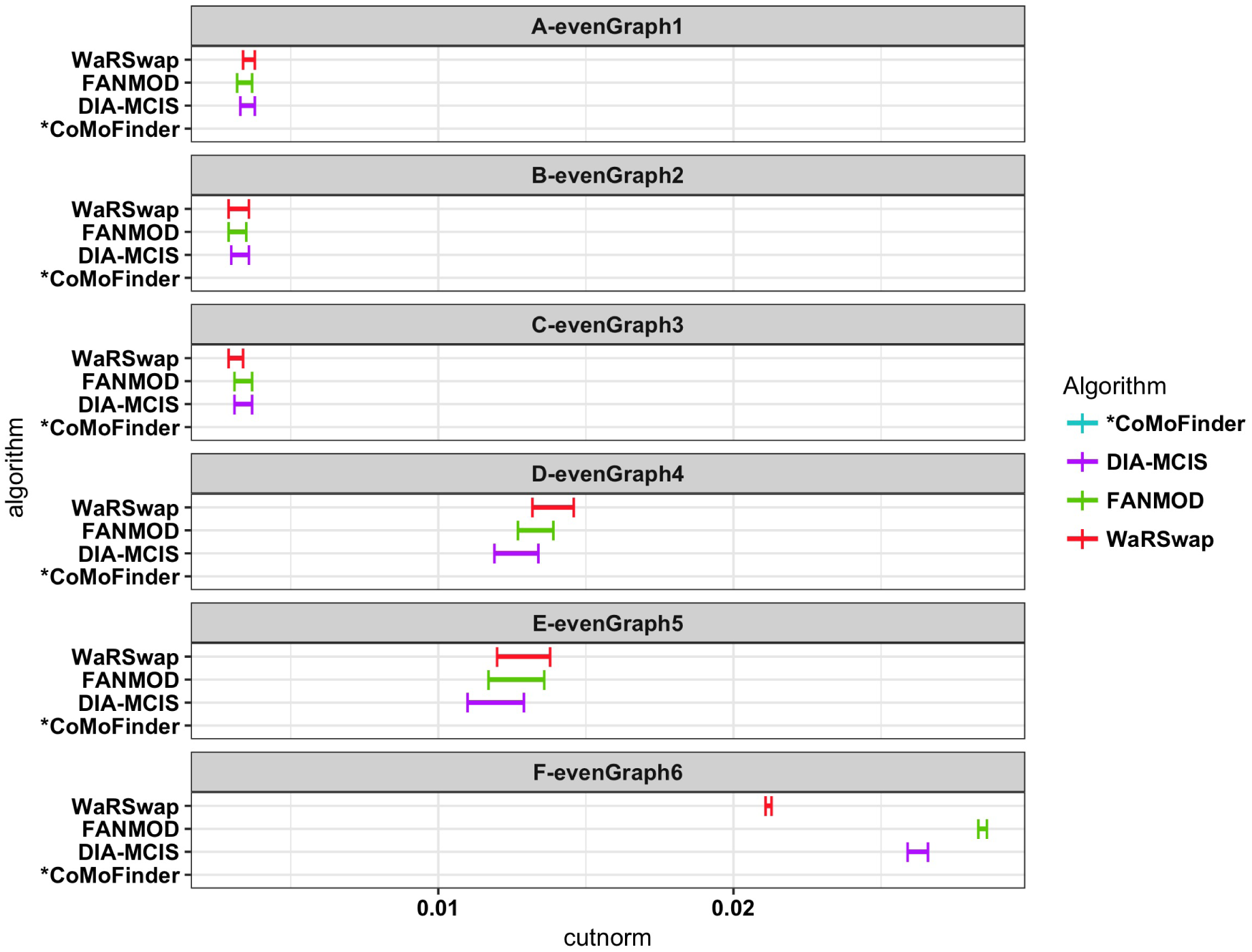
**Uniform/independent graph sampling performance evaluation on small even graphs.** For each small even graph and algorithm, 5000 graphs were generated. The cut norm estimates for each algorithm were computed using IndeCut. *The cut norm estimates for CoMoFinder were much larger than 0.06, therefore we removed CoMoFinder’s results from this figure for ease of comparison (see Supplementary Table S1 and Figure S4 for detailed results).

**Figure 4:**
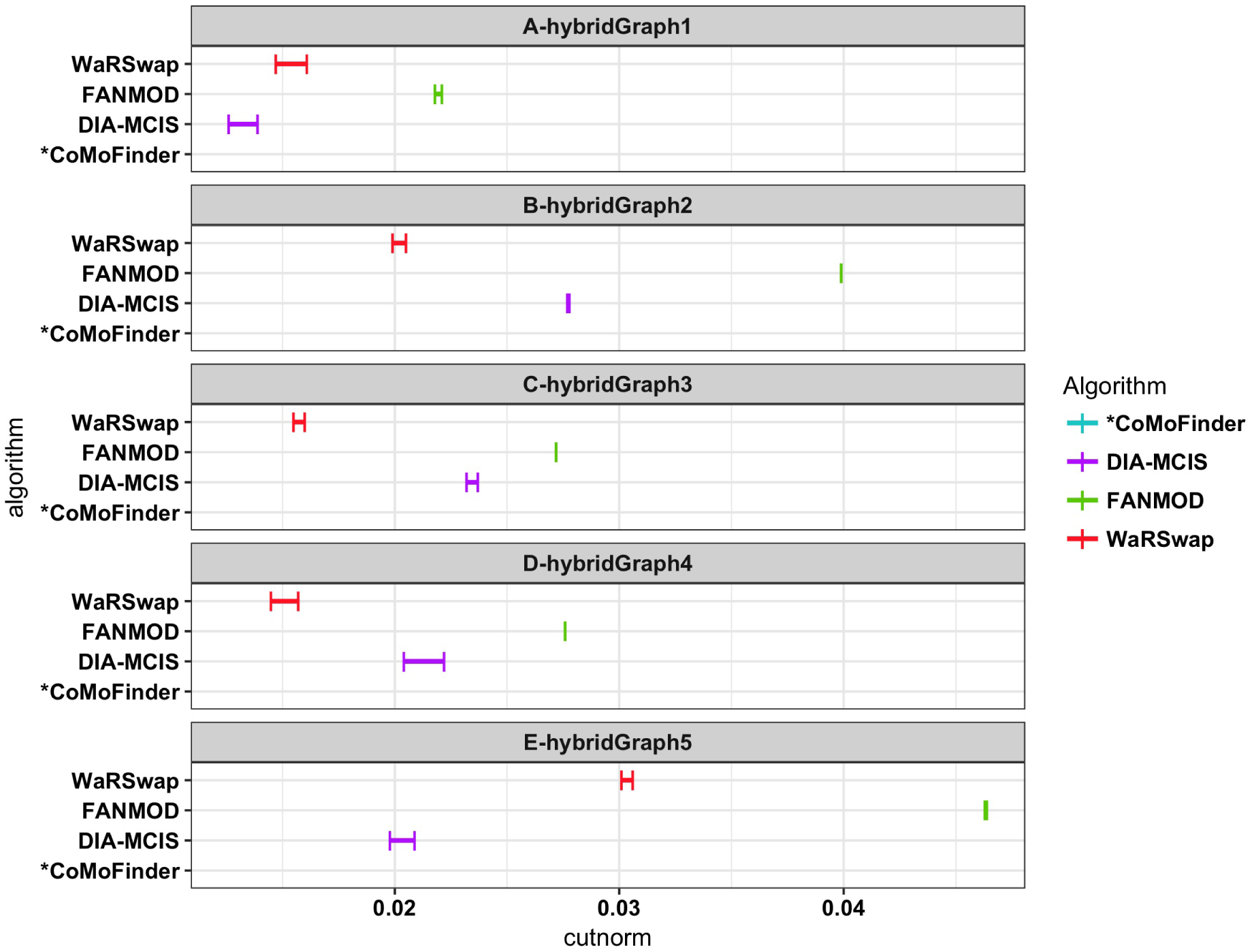
**Uniform/independent graph sampling performance evaluation on small hybrid graphs.** For each small even graph and algorithm, 5000 graphs were generated. The cut norm estimates for each algorithm were computed using IndeCut. *The cut norm estimates for CoMoFinder were much larger than 0.04, therefore we removed CoMoFinder’s results from this figure for ease of comparison (see Supplementary Table S1 and Figure S5 for detailed results).

In Fig 2, as the degree of unevenness for graphs increases from A to F, one observes that most of the example methods exhibit either increasingly poor sampling performance or relatively poor sampling performance in all cases. This is in contrast to the performance on the nearly regular graphs evaluated in Fig 3, where most of the methods have comparably strong performance. On the “hybrid” graphs in Fig 4, sampling performance varies widely among the methods. As has been discussed in previous network motif discovery studies, hub-containing graphs are highly problematic to many algorithms, whereas regular graphs are typically less troublesome for attaining uniform sampling performance ([3]). Fig 2 and Fig 4 confirm the fact that many algorithms have difficulty sampling from bipartite graphs containing large hubs, while most of the algorithms have comparably strong sampling performance when operating on a sample space of even or near-even graphs. The hybrid graphs highlight the necessity of IndeCut in determining the performance of each algorithm, particularly when the degree sequence of a graph yields no intuition with regard to the anticipated performance of any given method (Fig 4). We conclude that different graph topologies can produce vast performance differences when using the same algorithm, and that there are also large performance differences between algorithms on each graph topology.

#### 2.2.2 Real-world biological networks

In order to understand how these topologies interact in real biological graphs of interest, we examined two published biological networks with different degree sequences and scales: 1) An E-coli regulatory network that is a well-known example of a medium-size network (≈400 nodes and ≈600 edges) with a mixed degree sequence. This network contains two node types: transcription factors (TFs) and protein-coding genes (Genes). This network has been used as a case study by several network motif discovery studies including those published in conjunction with the FANMOD and CoMoFinder programs ([30, 32]). 2) A large human regulatory network (≈15, 000 nodes and ≈150, 000 edges) containing three different node types (TFs, miRNAs, and protein-coding genes). This network contains TFs that are “master regulators,” thus there are large source hubs in this network. This network has been used as a case study in CoMoFinder’s publication ([32]). We use this network to examine how efficiently IndeCut can assess the performance of graph sampling algorithms on a large realistic multi-layer network for which the user cannot anticipate motif discovery algorithm performance.

Since IndeCut operates on bipartite graphs as an input, the regulatory interactions between node types were broken into their component bipartite graphs (TF → TF, TF → Gene in the Ecoli network and TF → TF, TF → Gene, TF → miRNA, miRNA → TF, and miRNA → Gene in the human network). Fig 5 and Fig 6 show the resulting cut norm estimates for each graph and algorithm. In both figures, 5000 randomized graphs were generated using each of the example methods on each sub-network, then cut norm bounds were computed using IndeCut. However, DIA-MCIS is not able to operate on graphs with more than 2,035 nodes, therefore we do not have cut norm estimates for this method on the two large component graphs in the human network (TF → Gene and miRNA → Gene). These results indicate that at least one algorithm for every topology was able to achieve strong performance (near-uniform sampling performance). However, some graphs caused major performance difficulties for one or more algorithms. This highlights the importance of evaluating the performance of network motif discovery algorithms on biological networks of interest, particularly when considering costly and time-consuming experimental validations.

**Figure 5:**
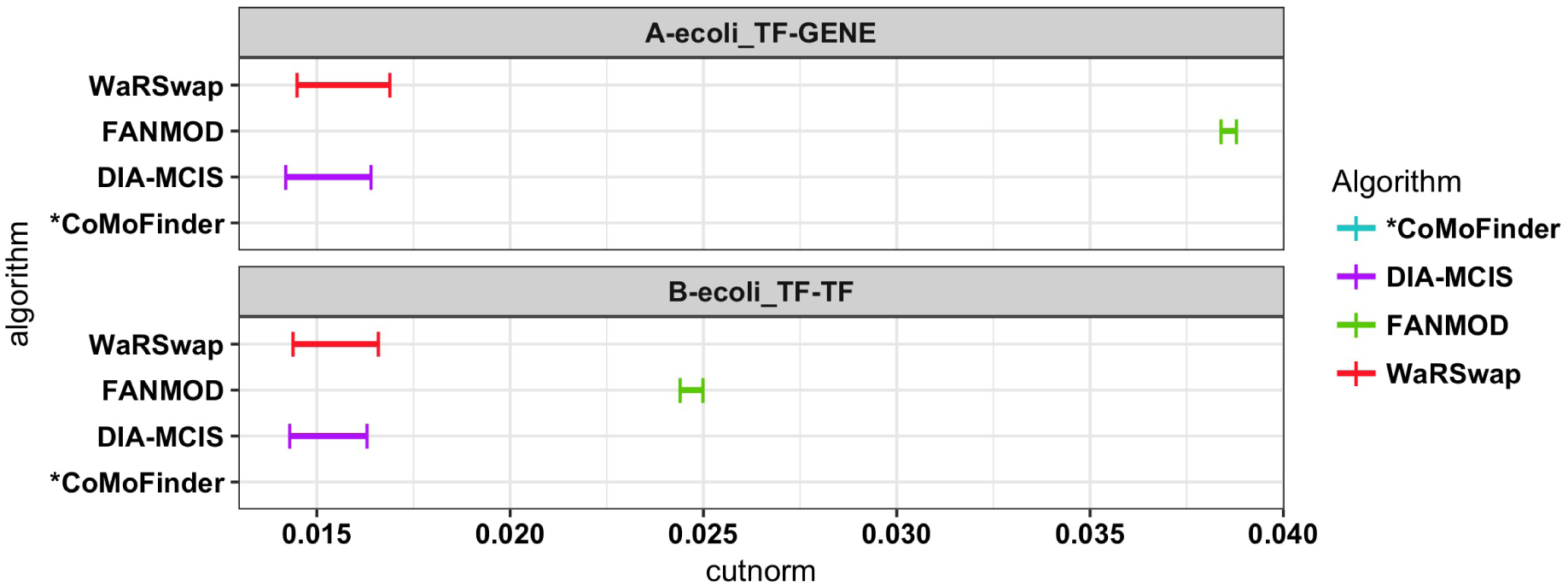
**Uniform/independent graph sampling performance evaluation on the Ecoli regulatory network.** For each small even graph and algorithm, 5000 graphs were generated. The cut norm estimates for each algorithm were computed using IndeCut. (A) Cut norm bounds resulting from running IndeCut on the Ecoli TF → TF network. (B) Cut norm bounds resulting from running IndeCut on the Ecoli TF → Gene network. *The cut norm estimates for CoMoFinder were much larger than 0.04, therefore we removed CoMoFinder’s results from this figure for ease of comparison (see Supplementary Table S1 and Figure S6 for detailed results).

**Figure 6:**
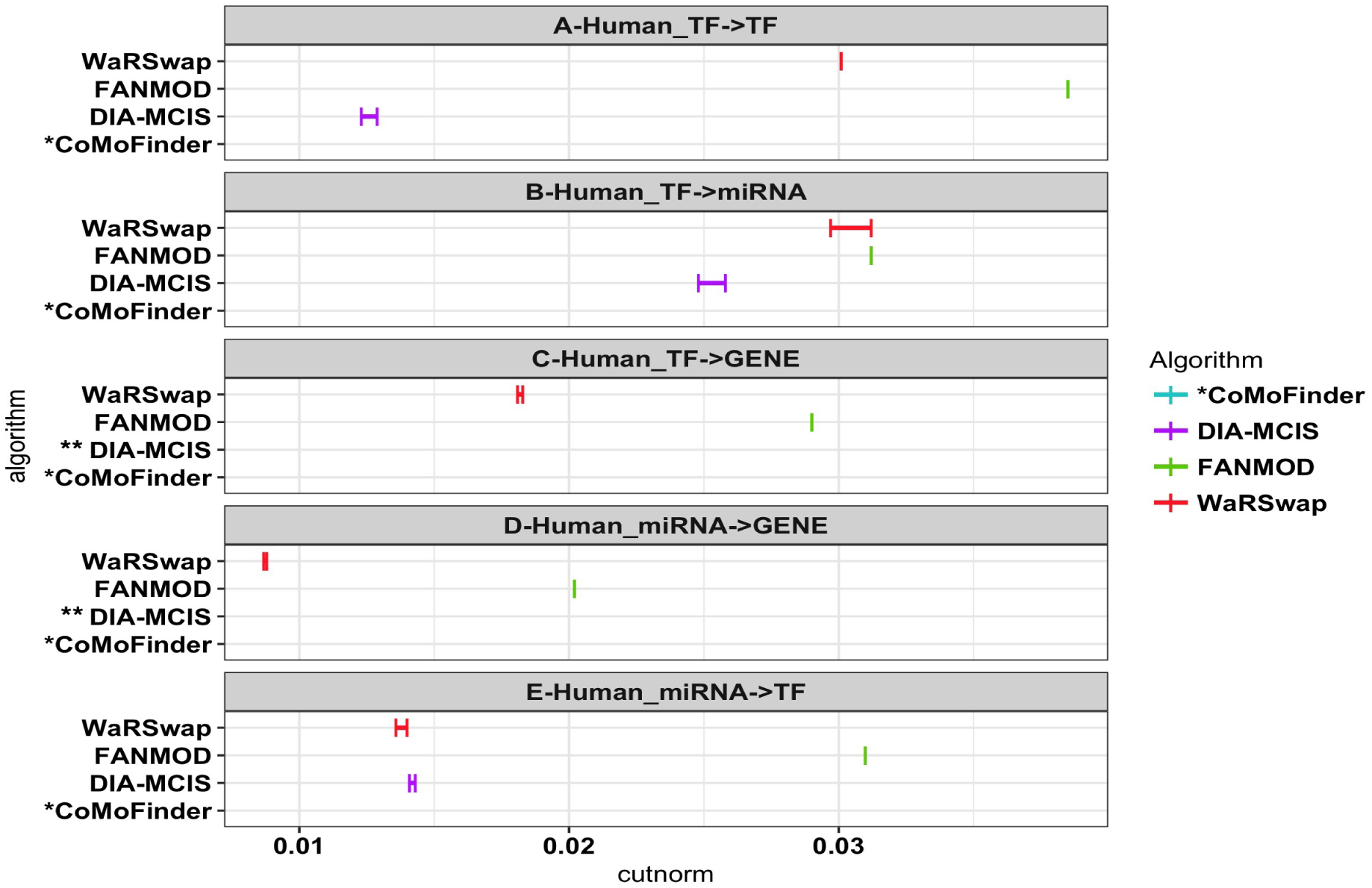
**Uniform/independent graph sampling performance evaluation on the Human TF-miRNA-Gene network.** For each small even graph and algorithm, 5000 graphs were generated. The cut norm estimates for each algorithm were computed using IndeCut. *The cut norm estimates for CoMoFinder were much larger than 0.04, therefore we removed CoMoFinder’s results from this figure for ease of comparison (see Supplementary Table S1 and Figure S7 for detailed results). **The cut norm estimates for DIA-MCIS are absent because this algorithm is not able to operate on graphs with more than 2,035 nodes.

**Table 1:** **Run time of** IndeCut **on a variety of graphs of different sizes.** Run time of IndeCut using a Linux machine with 48 Intel (2.40GHz) processors and 256GB of RAM.

| Graph | Number of nodes | Number of edges | *IndeCut* run-time |
| --- | --- | --- | --- |
| uniFanG1 | 11 | 20 | 10 s |
| hexaFanG1 | 66 | 120 | 48 s |
| regularG2 | 92 | 1058 | 1 min |
| Human TFTF | 174 | 644 | 3 min |
| Human TFmiR | 332 | 1,237 | 8 min |
| Human miRTF | 606 | 2,594 | 18 min |
| Human TFGENE | 9,055 | 25,748 | 4 days |
| Human miRGENE | 12,185 | 115,421 | 14 days |

IndeCut evaluates graphs on the order of several thousand nodes and tens of thousands of edges within a few minutes to a few days using standard hardware. Table 1 provides IndeCut’s observed run time on each graph and algorithm. To put these run times into perspective, network motif tools typically take several days simply to provide an output for graphs of this size, using a small number of iterations that does not guarantee meaningfully accurate performance (we discuss the number of iterations necessary for optimal performance for each sampling method in the next section). Using a commercial optimization package such as Guorbi or Mosek (in contrast to the open-source package CSDP that we use here) will result in speed improvements to IndeCut. Thus, considering time costs of running network motif finding algorithms themselves as well as the enormous potential laboratory costs of attempting to validate inaccurate results, IndeCut presents a very practical method for making an informed network motif discovery algorithm choice on biological networks of study.

### 2.3 IndeCut enables estimation of the number of samples necessary for an algorithm to achieve an accurate and reproducible outcome

For a very large network with hundreds of thousands of nodes and edges, running a network motif discovery program - even with the minimum number of samples recommended in the user manual - generally takes days to months, even when parallel processing is available. To date, there has been no method for evaluating the number of sample graphs necessary to produce a valid result (even if an algorithm is able to produce such a result). IndeCut is the first method to enable estimation of the number of sample graphs necessary to achieve the most accurate performance that each algorithm can provide. Intuitively, the larger the number of graphs sampled, the more accurately a program can “characterize” the nature of the entire background network sample space, leading to more accurate performance. Within a certain range of sample sizes, adding more graphs to a sample may result in a large performance increase. However, it is expected that beyond a certain sample size, performance increase per additional graph sampled will start to plateau (reach a point of diminishing returns). Here we use IndeCut to evaluate how the performance of a sampling algorithm improves as the number of graphs in a sample increases. We examine where a performance plateau occurred for each graph and algorithm. Furthermore, we provide an example from the literature that demonstrates the necessity of using IndeCut before performing any network motif discovery on a network of interest in real biological applications.

Given the space of all sampled graphs produced by an algorithm {*G*_1_, *…, G*_*n*_}, we generated *m* sets of samples {*S*_1_, *…, S*_*m*_} in which the set *S*_1_ contained the first 100 sample graphs {*G*_1_, *…, G*_100_}, set *S*_2_ contained all of the samples from *S*_1_ plus the next 100 samples {*G*_101_, *…, G*_200_}, and so on, until *S*_*m*_ contained all of the sample graphs {*G*_1_, *…, G*_*n*_}. We then used IndeCut to compute cut norm estimates for each set of subsamples *S*_*i*_, in order to identify an approximate sample size at which the cut norm estimate for *S*_*i*_ became very close to the cut norm estimate for the entire sample space *S*_*m*_ = {*G*_1_, *…, G*_*n*_}. Fig. 7 shows a visualization of the relationship between the number of samples and the cut norm estimates for a large biological network (TF → Gene network extracted from the Human regulatory network). These results show that overall, the approximate number of samples that is needed to approach best possible performance for each algorithm varies based on the topological features of the examined graph. For some algorithms, good performance is achievable even on topologies with extremely uneven degree sequences, but as expected, in all cases this comes at a cost of an increased number of samples. On the other hand, for more even degree sequences, a large number of samples is sometimes unnecessary (see Supplementary Figure S10, S11, and S12).

**Figure 7:**
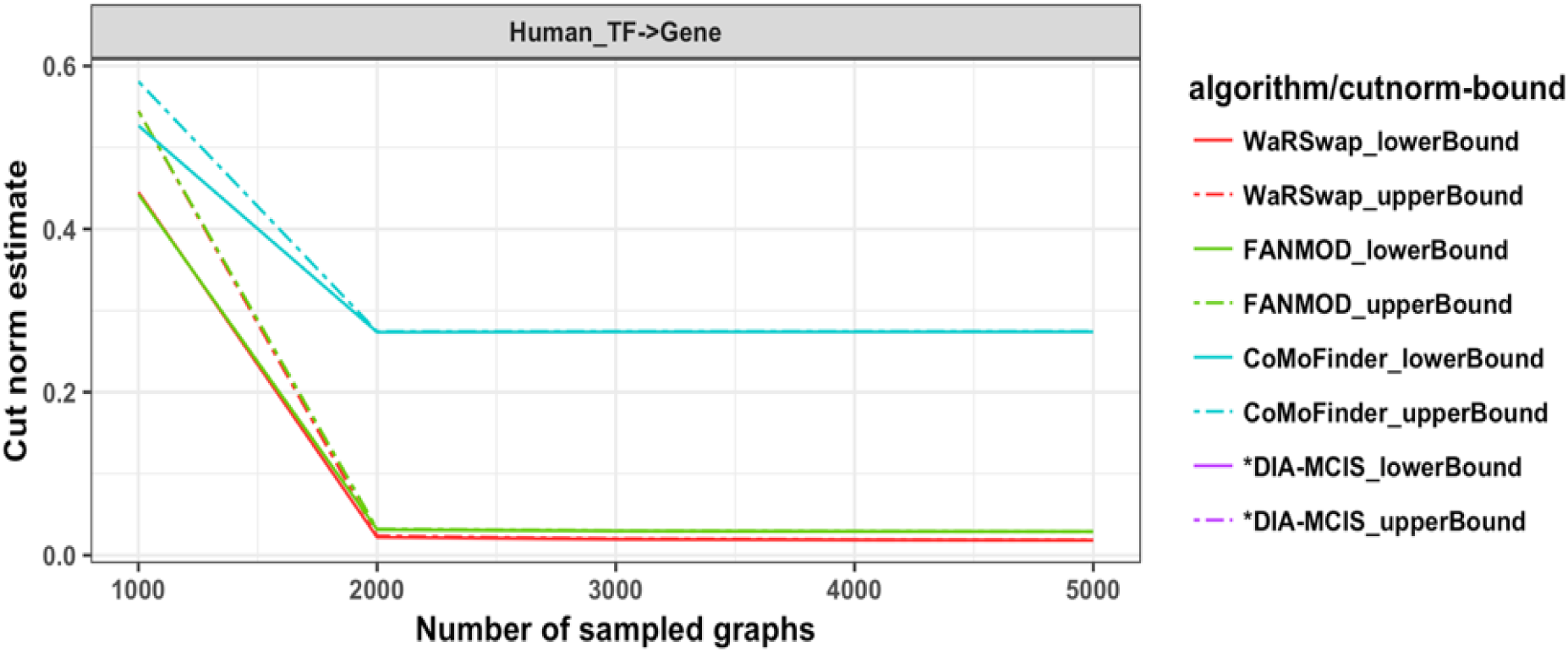
**Relationship between the number of samples vs. sampling performance for Hu-man TF** → **Gene network.** All 5000 samples previously generated by each algorithm for the Human TF → Gene network were collected and subsampled into five sets (1000, 2000, *…*, 5000 samples in each set, respectively). IndeCut was used to compute the cut norm estimates (lower and upper bounds) for each set of samples and algorithms. Cut norm values closer to zero represent a more uniform/independent sampling. This network has 9,055 nodes and 25,748 edges. *The cut norm estimates for DIA-MCIS are absent because this algorithm is not able to operate on networks with more than 2,035 nodes.

Our results show that IndeCut can help users to make an informed choice of the number of samples for each network and algorithm. In order to demonstrate how the reproducibility of a network motif outcome can be problematic without using a method such as IndeCut, we selected a published work ([33]) that reports network motifs in a Drosophila regulatory network. The authors have used FANMOD ([30]) to detect enriched 3-node network motifs in 100 sampled networks. To examine the reproducibility of the reported motifs, we ran FANMOD on the original network for 50 times, where for each time, 5000 samples were generated and the significance of 3-node subgraphs was computed for different subsets of samples (100, 200, 300, *…*, 5000 samples). Those 3-node subgraphs with p-value less than 0.01 and Z-score greater than 2.0 were considered in our analysis to be network motifs. In our results, motif5 in Fig7B of ([33]) (motif c in Fig. 8) was never observed in any iteration. It is likely that the relatively rarity of this subgraph in the portion of the sample space of randomized graphs that is explored by FANMOD at 100 iterations makes the single sample tested in ([33]) highly vulnerable to spurious observation. We also observed two significant network motifs, both of which were missed in the original work. Fig. 8 shows that with more than 1000 samples these two motifs are detected consistently, but with a smaller number of samples they do not reliably appear.

**Figure 8:**
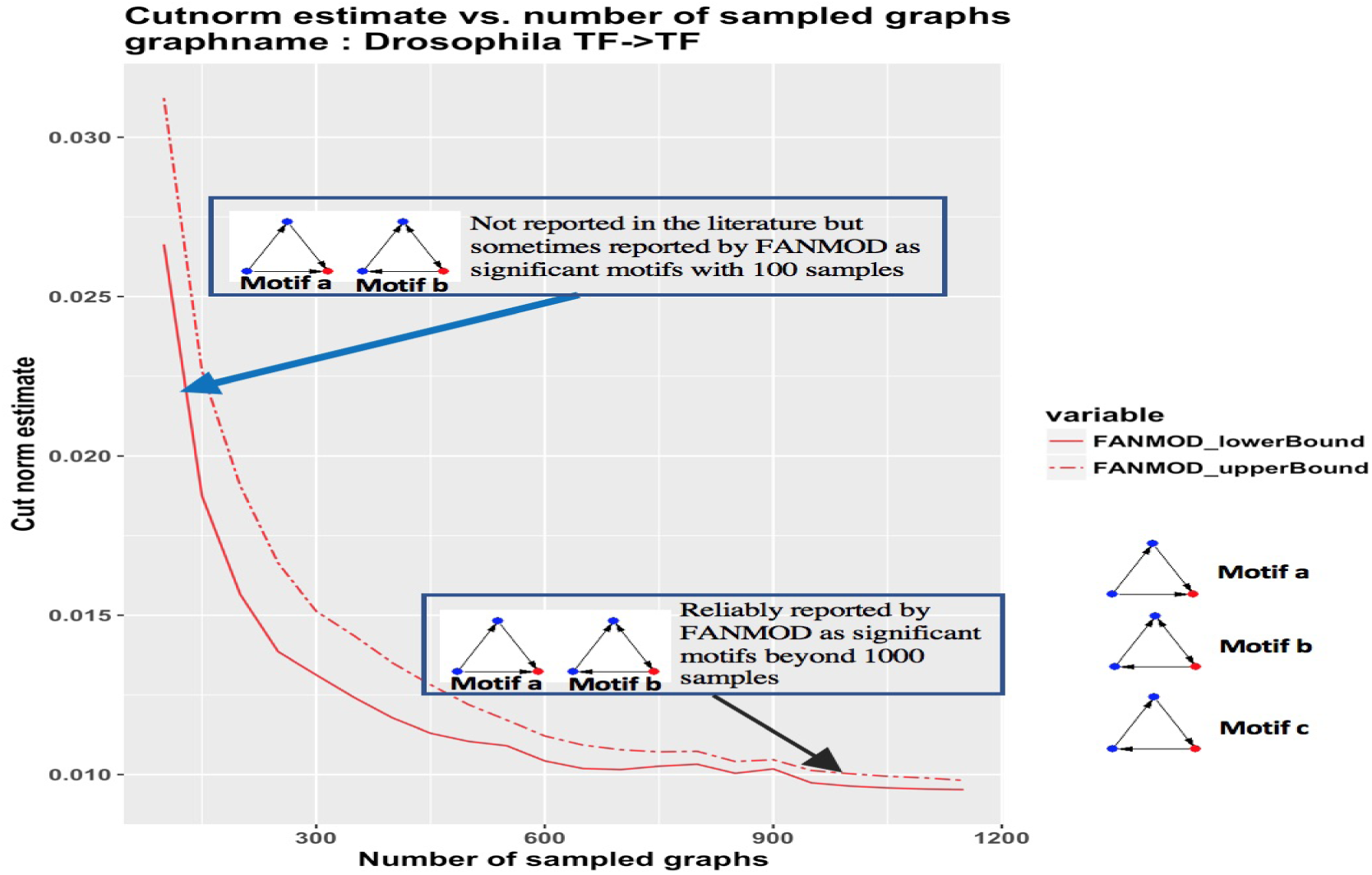
**The relationship between cut norm estimates, number of samples and network motif outcome on Drosophila network.** FANMOD was run on the Drosophila network ([33]) for 50 iterations. Motifs *a* and *b* are not reported in ([33]). These motifs were both found to be significant in a small proportion of trials at 100 sampled graphs (blue arrow). Motifs *a* and *b* are reliably detected beyond 1000 samples. Motif *c* was reported in ([33]) but was never observed as significant in any trials.

We used IndeCut to compute the relationship between the number of samples and the sampling performance of FANMOD on this network (Fig. 8). The performance plot in Fig. 8 shows that even for a moderately sized and relatively even graph such as the TF → TF layer extracted from the original network, at least ≈1000 samples are required to reach a performance that is close to the best possible performance of the algorithm. However, taking only 100 samples in the original analysis ([33]) is disturbingly vulnerable to reporting motifs that are artifacts of insufficient sampling, as well as to missing highly significant network motifs. It is completely understandable that given the long run-times required by many motif finding software implementations and no guidance on sufficient sampling, a relatively small number of samples was chosen for this analysis. IndeCut provides a practical solution for researchers using network motif discovery packages. Even if the initial package chosen is deemed inappropriate for accuracy given run-time, an alternative package or parallelized algorithm version can be evaluated.

Our results show that making an informed choice for the number of background sample graphs required for each algorithm and input network is incredibly important. In large real-world biological networks, we observe that even a “blanket policy” of generating a large number of graphs may not achieve reasonable performance for a given algorithm and graph topology (For example, FANMOD in its user manual recommends ≈1000 samples whereas Fig. 7 shows that at least 2000 samples are required to achieve reasonable performance in this case). We demonstrated that even for small to medium sized networks, understanding the appropriate number of samples is essential to attaining an accurate and reproducible outcome for each network and algorithm.

### 2.4 IndeCut elucidates when and why performance differences arise among network motif finding algorithms on different graph topologies

As the first method to evaluate performance of network motif finding strategies, IndeCut directly enables a new evidence-based framework for understanding when and why performance challenges occur. To demonstrate how IndeCut helps to achieve this goal, here we use the performance outcomes from IndeCut on our set of examined graphs from this study to analyze why certain graph topologies have been historically challenging for some classes of algorithms. We show that in cases of graphs with uneven degree distributions characteristic of biological networks, network motif discovery algorithms based on the graph randomization strategy known as “edge-switching” —the default strategy for the most commonly used algorithm —is vulnerable to highly non-uniform and/or non-independent sampling; thus this strategy is prone to spurious results on these networks. We use the concept of an edge-switching graph to show why this is the case. In essence, edge-switching algorithms produce a sampling bias by spending too much time sampling graphs that can be reached from the starting graph via a small number of edge-switches.

We first develop a new concept for visualizing how each sampling strategy behaves when sampling from the space of all possible graphs associated with a given degree sequence. We consider here networks that are small enough that the sample space of all bipartite graphs with fixed degree sequences is of reasonable size. Given a sample space containing *n* graphs, we create an “edge switch graph” (ESG) of *n* nodes where each node represents an individual graph in the sample space {*G*_1_, *…, G*_*n*_}. In the ESG, an undirected edge connects two nodes (graphs *G*_*i*_ and *G*_*j*_) when exactly one edge switching operation converts *G*_*i*_ to *G*_*j*_ and vice versa. Fig. 9 demonstrates the construction of a 5-node ESG graph given a degree sequence of R = C = {2,1,1}. We then applied a graph clustering algorithm ([34]) to the resulting ESG to detect clusters of closely connected graphs. This clustering method maximizes the number of paths within a cluster and minimizes the number of paths between clusters. Therefore, each cluster represents a collection of graphs that are close to each other in the sense that a relatively small number of edge-switching operations is needed to convert one to the other. These edge switch graphs allow us to visualize the topology of the space of all bipartite graphs with fixed degree sequence.

**Figure 9:**
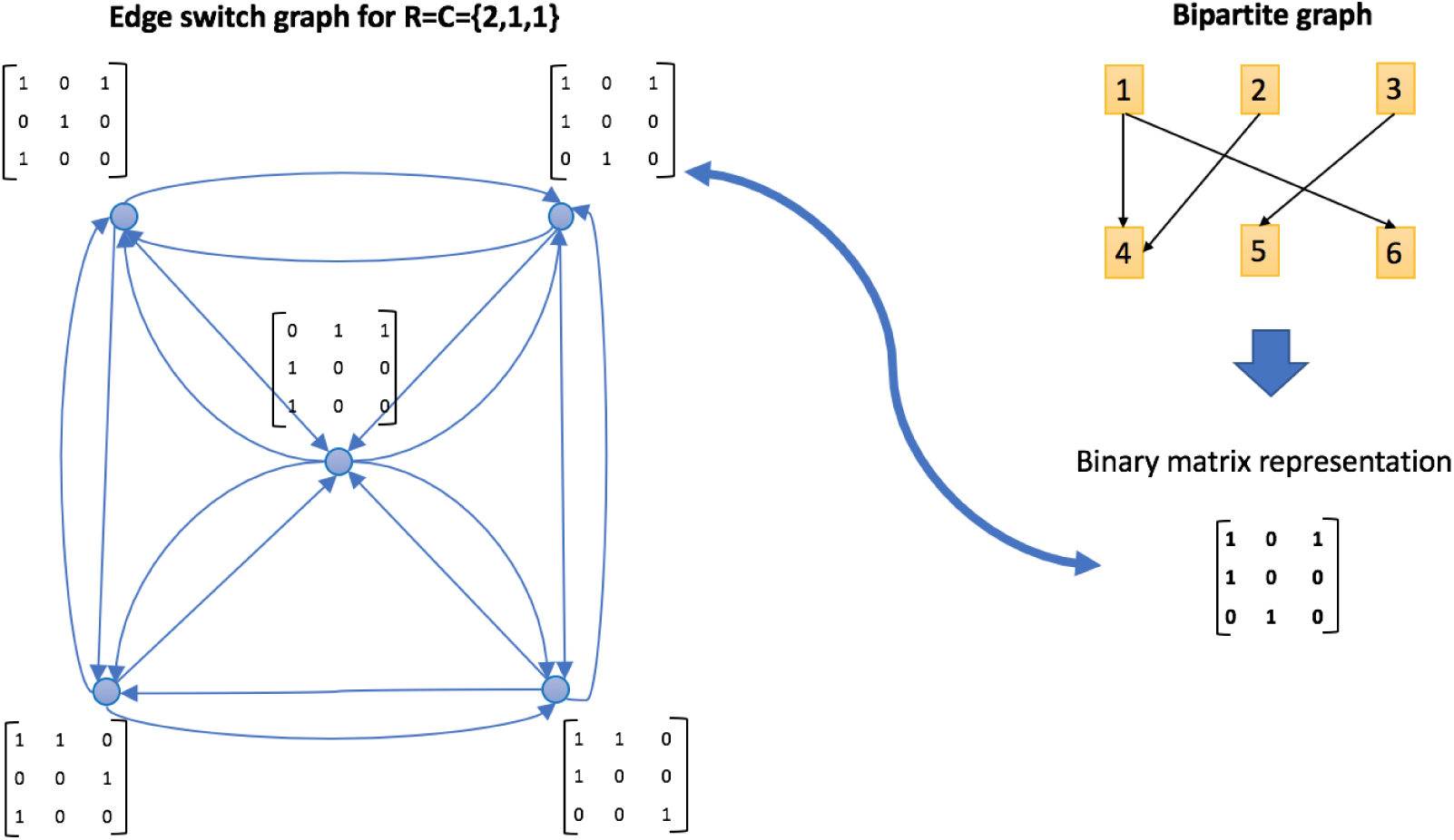
**Constructing an ESG graph.** The bipartite graph (top right) with degree sequence of R=C= {2,1,1} produces a sample space containing five different graphs which are represented as nodes in the ESG (left). The zero-one matrices represent the edge configuration of each node. An edge connects two nodes (graphs) which can be converted to each other by preforming one edge switch.

Fig. 10B shows an ESG graph constructed from an in-degree sequence of *R* = {2, 1, 1, 2, 1, 1} and an out-degree sequence of *C* = {2, 1, 1, 2, 1, 1}. The sample space of this graph has 5400 elements. After running the graph clustering algorithm, 10 separate clusters were detected. We executed each algorithm on the given degree sequence to produce 10,000 sample graphs per algorithm. We then calculated the number of times each algorithm returned a graph falling within each of the clusters (normalized by cluster size). This indicates how each of the examined algorithms samples its space with respect to these clusters (an equal number of graphs sampled within each cluster indicates a more uniform sampling method). Fig. 10C-F shows “cluster-time” diagrams, visualizing how much time each algorithm has spent in each cluster. In a cluster-time diagram, nodes represent clusters (in the ESG), and the size of each node represents the fraction of graphs sampled in the corresponding cluster compared to all graphs sampled (i.e. the total “time” the algorithm spends in a given cluster). The larger a node appears, the more time that has been spent sampling from the associated cluster by a given algorithm. A method that samples uniformly will result in a cluster-time diagram with precisely equal node sizes. As a measure of this outcome, we take the fraction of time spent in each cluster and compute the entropy of this collection of fractions to summarize the “evenness” of the sampling regime (larger numbers are better). The entropy value for each algorithm is noted above the corresponding cluster-time diagram in Fig. 10C-F (See Supplementary FigureS8,S9 for more examples).

**Figure 10:**
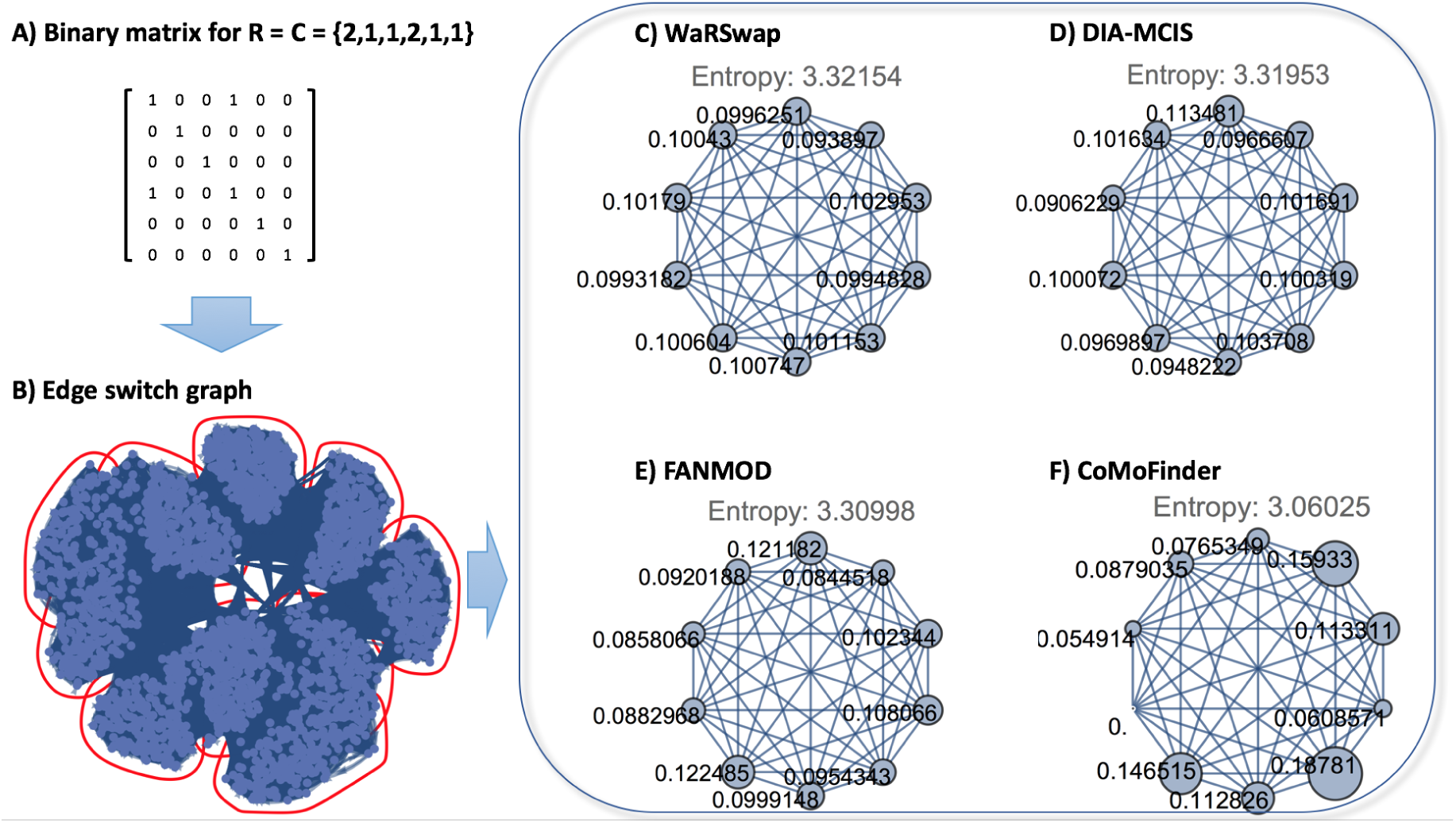
An example ESG graph for degree sequence R=C={2,1,1,2,1,1}. A) The 0-1 matrix of the initial graph. B) ESG graph corresponding to entire sample space of given degree sequence (blue dots represent graphs in sample space) with detected clusters (in red). C-F) Cluster-time diagrams for each examined algorithm; nodes represent clusters in the ESG with node size indicating the fraction of times a given algorithm sampled graphs in that cluster.

Algorithms based on edge switching as a method of randomized background network generation, such as FANMOD and CoMoFinder, generally spend a substantially uneven amount of time in different clusters. Intuitively, this is the case because an edge switching method begins with the input graph (the original biological network) and then performs a series of edge-switching operations, resulting in one background graph in the sample. Each series of operations corresponds to a path in the ESG. In general, real biological networks have sample spaces in which some graphs in the space are “easy” to reach via edge-switching operations from the initial “real” network, while others are more “difficult” to reach from this initial graph in the sense that one must select a rare sequence of edge switches in order to reach these graphs. FANMOD’s strategy selects sequences of edge-switching operations without any condition on the number of times that the same pair of edges can be selected for an endpoint switch. CoMoFinder also selects sequences of edge-switching operations, but disallows revisitation of the same pair of edges. Effectively, when traversing a path in an ESG away from the initial graph using CoMoFinder’s strategy, the number of paths available to reach a destination graph from the current state is limited as compared to FANMOD’s strategy.

In contrast, WaRSwap and DIA-MCIS do not use the edge-switching method, but rather generate each sample graph by placing edges between source and target nodes using a weighted sampling scheme (thus there is no direct relationship between a sampled background graph and the initial input graph). In Fig. 10, the size of the cluster nodes is nearly even for WaRSwap and DIA-MCIS, and the corresponding entropy values are higher as compared to FANMOD and CoMoFinder. This is generally true on graphs examined throughout our results. However, there are also performance differences between DIA-MCIS and WaRSwap on large hub-containing graphs (Fig. 2). Depending on the weighted sampling strategy, these algorithms can behave differently while sampling from graphs containing large hub nodes. As shown in Fig. 10, WaRSwap exhibits nearly equal cluster-time as compared to DIA-MCIS. This can be explained by the fact that WaRSwap uses a dynamically weighted sampling formula to distribute edges between source and target nodes according to their current target degrees at each stage of edge placement. WaRSwap’s weighting formula helps to correct the strong tendency of source hub nodes (nodes with a higher out-degree) to connect to target hub nodes (nodes with a higher in-degrees), a phenomenon that leads to oversampling of hub-hub connections. While DIA-MCIS is generally able to perform well on all topologies, in the case of certain highly uneven graphs, a static weighting strategy is susceptible to undersampling of rare graphs (those with few/no hub-hub connections). Overall, results indicate that either of these strategies would provide a safer choice than an edge-switching strategy on biological networks.

## 3 MATERIALS AND METHODS

### 3.1 Definitions and overview

A graph *G* = (*V, E*) is an object describing the relationships between a set of objects *V*, referred to as the *vertex set*, by a set of directed *edges* (*v*_*i*_, *v*_*j*_) *∈ E* where *v*_*i*_, *v*_*j*_ *∈ V*. In this work, we define a network as a two-layered or *bipartite* graph *G* containing *m* source nodes {*S*_1_,…, *S*_*m*_} and *n* target nodes {*T*_1_,‥, *T*_*n*_} where a single directed edge connects a source node to a target node. The number of edges coming into a node is called its *in-degree* and the number of edges coming out from a node is called its *out-degree*. In a bipartite graph *G*, source nodes and target nodes have zero in-degrees and zero out-degrees, respectively (edges emanate only from source nodes, and are directed only to target nodes). The structure or topology of a graph *G* can be described by its in-degree and out-degree sequences.

A bipartite graph *G* can be represented as a binary matrix *A ∈* {0, 1}^*m×n*^ where *m* is the number of source nodes (number of rows) and *n* is the number of target nodes (number of columns). *A*_*i,j*_ = 1 means that there is a direct edge from *S*_*i*_ to *T*_*j*_, and *A*_*i,j*_ = 0 means there is no edge between them. The row sums *R* = (*r*_1_, *…, r*_*m*_) and column sums *C* = (*c*_1_, *…, c*_*n*_) of matrix *A* represent the out-degree and in-degree sequences of *G*, respectively. Collectively, they are referred to as the degree sequences of a graph. Hence, we have that Σ_*i*_ *A*_*i,j*_ = *C*_*j*_ and Σ_*j*_ *A*_*i,j*_ = *R*_*i*_.

### 3.2 Proof of correctness of IndeCut

For a given degree sequence (*R* and *C*) of a bipartite graph *G*, the sample space of graph *G* defines a set of all distinct graphs that can be constructed from *R* and *C*. In order to understand whether a set of graphs generated by a randomization strategy are selected uniformly and independently from the sample space, we must compute an expected incident matrix. We define *E* as the expected/average matrix over all possible graphs producible from *R* and *C*. Given *N* random graphs 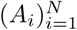 generated by a network motif finding algorithm from *R* and *C*, we would like to compute the distance of the sample average of these *N* matrices from *E* as follows.

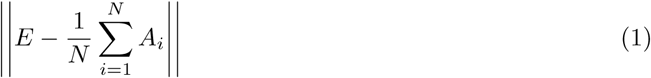

Performance of uniform and independent sampling by an algorithm can be verified if this difference is small (in probability) for some large *N* and some appropriately defined matrix norm ∥.∥ However, this approach is not feasible since the matrix *E* is extremely difficult to calculate. Except in very special cases of *R* and *C*, the only known way to compute *E* would be to enumerate all bipartite graphs having degree sequences *R* and *C*, form their incidence matrices, and average them. Since the sample space of large biological graphs is enormous, there is no practical computational method for producing all of the incidence matrices; accordingly, calculating *E* is currently intractable. Thus, we take an alternative approach as follows: we utilize results from [29] wherein a matrix *Z*, called “Maximum Entropy Matrix”, is obtained in a computationally tractable manner that can be thought of as a proxy for the average incidence matrix *E*. The *cut norm* ∥ *·* ∥_*C*_ is the appropriate matrix norm to use in this case.

### 3.3 Maximum Entropy Matrix computation

Let Σ(*R, C*) be the set of all binary matrices with row-sums *R* = (*r*_1_, *…, r*_*m*_) *∈*ℕ^*m*^ and column-sums *C* = (*c*_1_, *…, c*_*n*_) *∈*ℕ^*n*^. Throughout, we only consider *R* and *C* such that for every choice of 1 *≤ i ≤ m* and 1 ≤ *j* ≤ *n*, there exist at least two matrices *L, M* ∈ Σ(*R, C*) such that *L*_*i,j*_ = 0 and *M*_*i,j*_ = 1. This condition requires the space Σ(*R, C*) to be reasonably large.

We now recount pertinent theorems from [29]. The first gives an estimate of the number of bipartite graphs with degree sequences *R* and *C*: |Σ(*R, C*)|.

#### Theorem 1

([29, Theorem 1.1]). *Let*

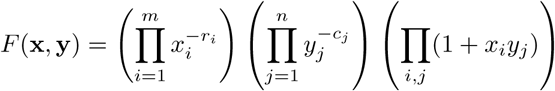

*for* x = (*x*_1_;…, *x*_m_) *and* y = (*y*_1_;…, *y*_n_). *Let*

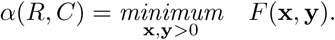

*Then*

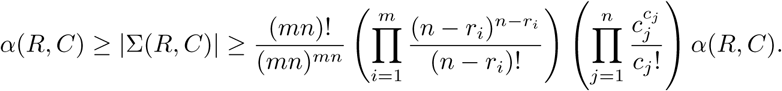

Taking the logarithm of *F* (**x**, **y**) gives a convex function on ℝ^*m×n*^, so *α*(*R, C*) may be efficiently computed. This allows us to define the *maximum entropy matrix*:

#### Definition 1

(Maximum Entropy Matrix). *Let* **x**^*\**^ *and* **y**^*\**^ *be the vectors that obtain optimality in the definition of α*(*R, C*). *Define Z ∈*ℝ^*m×n*^ *as*

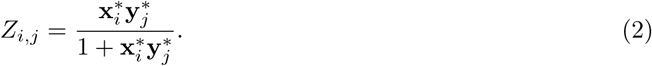

The maximum entropy matrix *Z* should be thought of as a proxy (in the cut norm, see Definition 2) for the average matrix of Σ(*R, C*) (see [29, Theorem 1.5]).

Let 𝒜 represent a given motif finding algorithm (thought of as a binary matrix valued random variable). Let 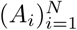 be *N* iterates of this algorithm and define

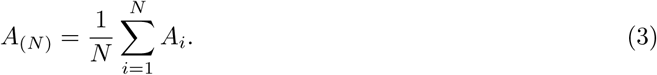

If the sequence (*A*_*i*_)_*i≥*1_ is a realization of a sequence of i.i.d. (independent and identically distributed) random matrices, then a consequence of [29, Theorem 1.4] is that with high probability, the norm ∥*Z - A*_(*N*)_∥_*C*_ is small. We can thus use

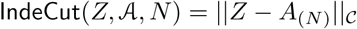

as a characterization of the performance of a motif finding algorithm A in terms of how well it samples the relevant space of bipartite graphs in an i.i.d. fashion: For large *N*, if one algorithm outputs matrices whose average is closer in the cut norm to *Z* than that of another algorithm, then the former algorithm’s heuristics sample the space Σ(*R, C*) in a more i.i.d. fashion.

We turn now to defining the cut norm and how to compute it.

### 3.4 Computing norms

It will turn out that the cut norm ∥ *·* ∥_*C*_ is difficult to compute (in fact, it is MAX SNP hard), but we will be able to relate it to another norm (∥ *·* ∥_*∞→*1_) that can be approximated with a semidefinite relaxation. We then round the solution of the semidefinite relaxation to get an estimate of ∥ *·* ∥_*∞→*1_ and hence of ∥ *·* ∥_*C*_. We begin with definitions of the norms of interest.

#### Definition 2.

*Let A ∈*ℝ^*m×n*^. *Define the following norms by*

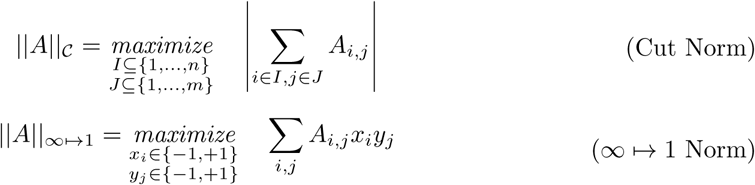

We denote the semidefinite relaxation of ∥*A*∥_*∞→*1_ by ∥*A*∥_SDR_:

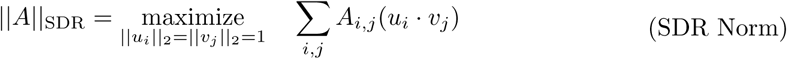

Note that ∥*A*∥_SDR_ can be converted to the following optimization problem:

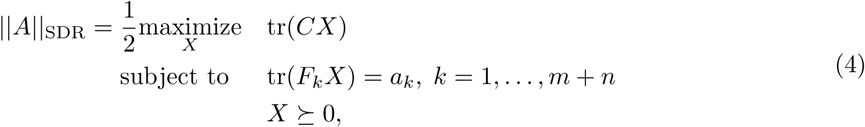

for

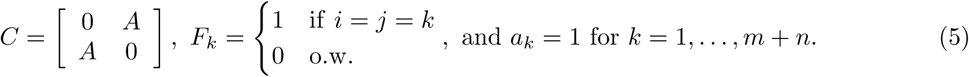

This form allows us to use popular computational packages to compute ∥·∥_SDR_. We utilize the computational package CSDP version 6.1.0 [35].

It turns out that for the matrices of interest, the norms ∥·∥_*C*_ and ∥·∥_*∞→*1_ are equal up to a factor of 4. Indeed, note that both the maximum entropy matrix *Z* defined in equation (2), as well as the average of the output matrices *A*_(*N*)_ defined in equation (3) have row/column sums equal to *R* and *C*. That is, Σ_*i*_ *Z*_*i,j*_ = Σ_*i*_(*A*_*n*_)_*i,j*_ = *R*_*j*_ and Σ_*j*_ *Z*_*i,j*_ = Σ_*j*_ (*A*_*n*_)_*i,j*_ = *C*_*i*_, hence the matrix *Z* - *A*_(*N*)_ has zero row and column sum. This allows us to obtain the well-known [28, 36] relationship between the norms ∥ *·* ∥_*∞→*1_ and ∥ *·* ∥_*C*_.

#### Proposition 2.

*If the matrix A has zero row and column sums then* Σ_*i*_ *A*_*i,j*_ = Σ_*j*_ *A*_*i,j*_ = 0, *then* ∥*A*∥_*∞→*1_ = 4∥*A*∥_*C*_.

*Proof.* For *I ⊆* {1, *…, n*} and *J ⊆* {1, *…, m*} the sets achieving the maximum in the definition of ∥*A*∥_*C*_, define *x*_*i*_ = 1 for *i ∈ I, x*_*i*_ = *-*1 for *i ∉I* and *y*_*j*_ = 1 for *j ∈ J, y*_*j*_ = *-*1 for *j ∉J*. Then

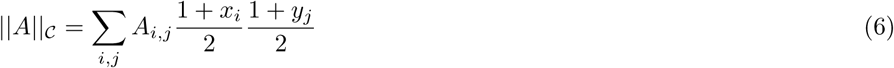

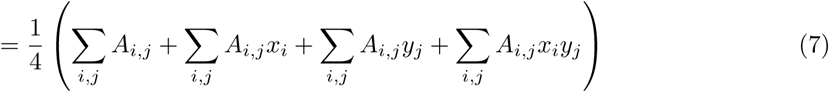

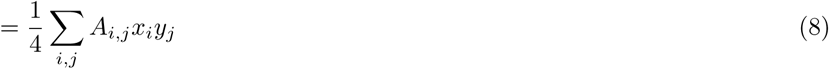

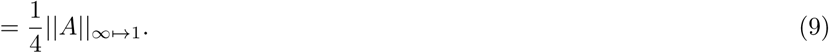

We now turn our attention to computing the cut norm ∥ *·* ∥_*C*_.

#### 3.4.1 Computing the cut norm ∥ *·* ∥_*C*_

Unfortunately, computing the cut norm directly is MAX SNP hard [28]. However, it is possible to obtain upper and lower bounds on the cut norm of a given input matrix. We use these bounds in the following way: for input matrices *A, B*, say there exist bounds *a*_*u*_, *a*_*l*_ and *b*_*u*_, *b*_*l*_ such that *a*_*l*_ *≤* ∥*A*∥_*C*_ *≤ a*_*u*_ and *b*_*l*_ *≤* ∥*B*∥_*C*_ *≤ b*_*u*_. If *a*_*u*_ *< b*_*l*_ or *b*_*u*_ *< a*_*l*_ then we may say unequivocally that ∥*A*∥_*C*_ *<* ∥*B*∥_*C*_ or ∥*B*∥_*C*_ *<* ∥*A*∥_*C*_ respectively. However, if the bounds overlap (*a*_*u*_ *≥ b*_*l*_ or *b*_*u*_ *≥ a*_*l*_) then no claim can be made about the relationship of ∥*A*∥ _*C*_ and ∥*B*∥ _*C*_.

In [28, Section 5.1], an algorithm was presented that computes such bounds. We use a slight modification of this algorithm that gives tighter bounds in practice as follows:

Given a matrix *A*, let *u*_*i*_, *v*_*j*_ *∈*ℝ^*m*+*n*^, for *i* = 1, *…, m, j* = 1, *…, n* be the optimal vectors obtained from the computation of ∥*A*∥_SDR_. Let *g*_*i*_ *∼*𝒩 (0, 1), *i* = 1, *…, m* + *n*, be independent standard normal random variables and let *G* = (*g*_1_, *…, g*_*m*+*n*_). Let

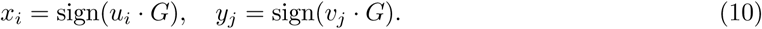

Now Σ_*i,j*_ *A*_*i,j*_*x*_*i*_*y j ≤* ∥*A*∥_*∞→*1_ since ∥*A*∥_inf*1→*1_ is the maximum value. However, there is a positive probability that 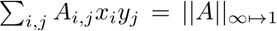. To observe this fact, let 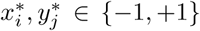 be such that 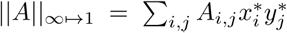. We can find at least one vector *G*^*\**^ such that 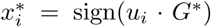 and 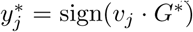 since this reduces to solving a solvable system of linear inequalities due to the *u*_*i*_, *v*_*j*_ being obtained from eigenvectors of the spectral factorization of *X* in the optimization procedure (4). Given such a *G*^*\**^, note that for any *a ∈*ℝ, *a >* 0, 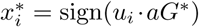 and 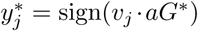. Hence, with probability at least 2^*-m-n*^, a randomly chosen *G* will result in obtaining the optimal 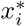 and 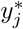. We do not attempt to make a more nuanced estimation of this probability since only bounds are necessary for our purposes.

Repeating the above rounding procedure a number of times and taking the maximum result, we obtain Algorithm 1 which we use to compute the bounds on the cut norm of a matrix *A*. In practice, we take the number of iterates of Algorithm 1 to be 1, 000. Denote the output of this algorithm with 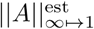. As a result, 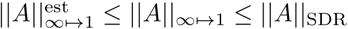, so combining these with proposition 2, we have the following estimation of the cut norm:

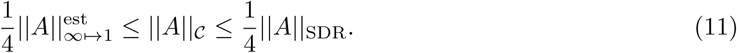

#### Algorithm 1

Cut norm lower bound

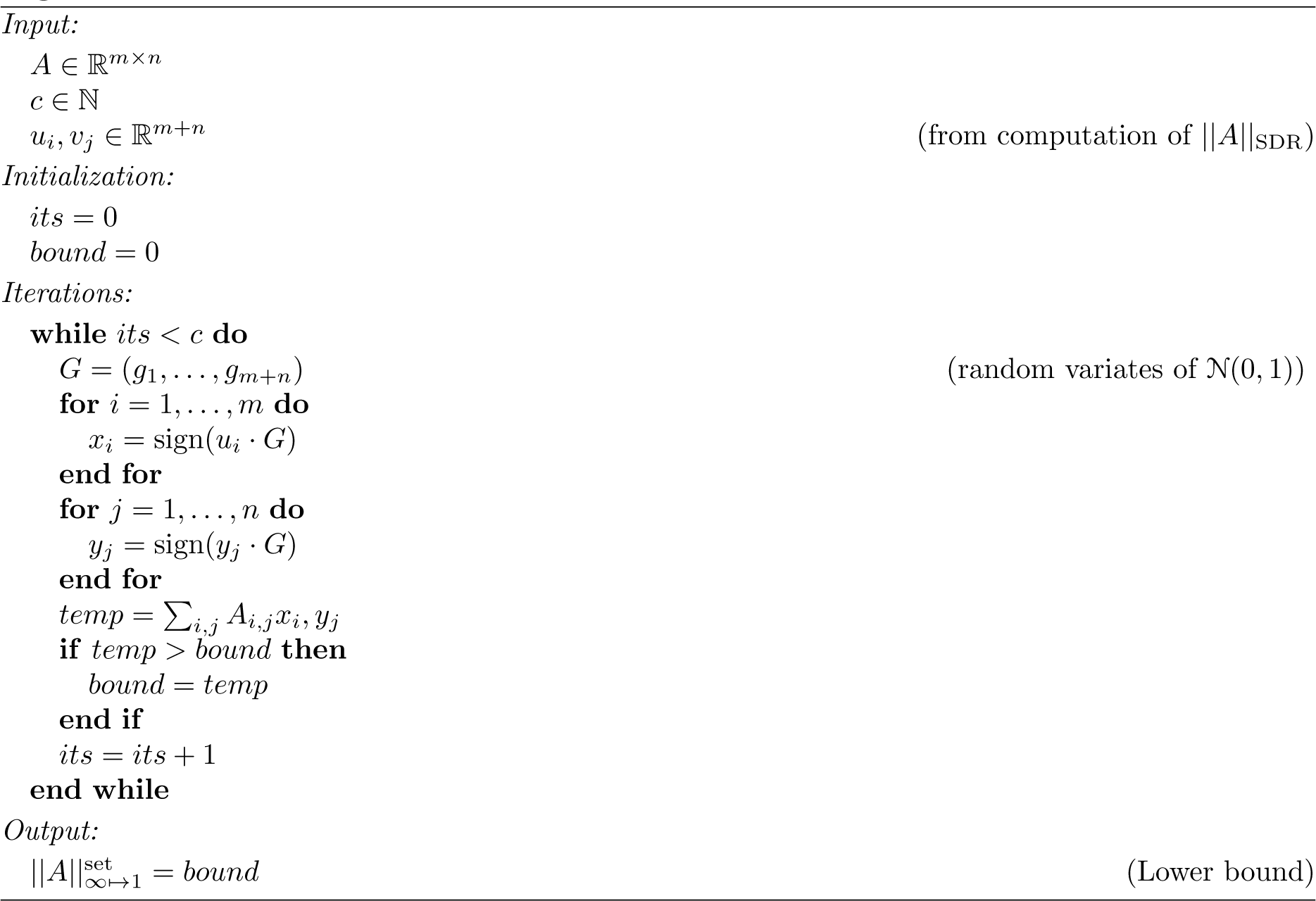

We apply this estimation to obtain bounds on the quantity of interest:

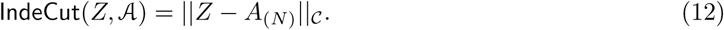

Now that we have the cut norm bounds for each network motif finding algorithm of interest, we can compare the performance of those algorithms as follows:

Let 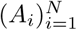 and 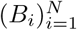be *N* random incidence matrices generated by two algorithms 𝒜 and ℬ. If for sufficiently large *N*, if we have

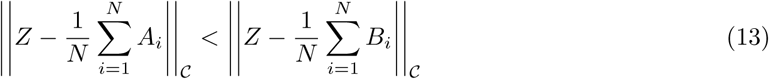

we can conclude that algorithm 𝒜 samples the space of all relevant bipartite graphs in a more i.i.d. fashion than algorithm *B*.

### 3.5 Networks and graphs

Two sets of graphs were created or selected for this study: 1) Manually constructed “toy” bipartite graphs with sizes ranging from tens of nodes to hundreds of nodes, representing different graph structures, including “even” or “near-even” graphs, “uneven” graphs, and “hybrid” combinations of in/out-degrees, and 2) Real biological networks.

*Real networks* - Two biological networks were obtained from literature and public databases. An *Ecoli* network representing a medium-size yeast transcriptional network was downloaded from [7]. This network contains two types of nodes: transcription factor (TF), and gene. Two layers of interactions (TF →gene, TF →TF) were extracted into separate bipartite graphs for application of IndeCut. A *Human* regulatory network was downloaded from http://encodenets.gersteinlab.org/, representing a network with thousands of nodes and edges. This network is used as a case study in the publication of CoMoFinder [32]. This network contains three types of nodes: TF, miRNA, and protein-coding gene. This network comprises five interaction layers: TF→TF, TF→miRNA, miRNA→TF, TF→gene, and miRNA→gene. Each of these layers forms a separate input bipartite graph for IndeCut.

See Supplementary Text1 for detailed description of each network motif discovery algorithm.

### 3.6 Description of edge switch graphs

We detail here how the edge switch graphs (ESG’s) were created. Given in and out-degrees *R* and *C*, we generate all possible bipartite graphs {*G*_1_, *…, G*_*N*_} with in/out-degrees *R* and *C*. The edge switch graph *G*_ESG_ is an undirected graph with vertex set *V* = {*G*_1_, *…, G*_*N*_} and edge set *E* defined as follows: for *G*_*i*_, *G*_*j*_ *∈ V*, the undirected edge (*G*_*i*_, *G*_*j*_) is an element of *E* if and only if the graph *G*_*j*_ can be obtained as a result of performing one edge switch on *G*_*i*_. In more detail, this means that the graphs *G*_*i*_ and *G*_*j*_ have the same vertex set, and identical edge sets, except for one pair of edges (*x, y*) and (*u, v*) present in the edge set of *G*_*i*_ but absent in the edge set of *G*_*j*_, and one pair of edges (*x, v*) and (*u, y*) present in the edge set of *G*_*j*_ but absent in the edge set of *G*_*i*_.

A graph clustering algorithm known as modularity clustering [34] was then applied to the edge switch graph *G*_ESG_ to identify clusters that maximize the number of within-cluster edges while minimizing the number of between-cluster edges. Let *L* be the number of clusters found.

Given a graph sampling algorithm 𝒜, the output of 𝒜 can be viewed as sampling vertices of the ESG. Define a count vector *count*^𝒜^ *∈*ℕ^*L*^ as a vector indexed by the clusters found above, with 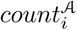 being equal to the number of times the algorithm 𝒜 returned a graph found in cluster *i*.

A “cluster-time” graph is then created with vertices corresponding to the clusters found above, and edges between two pairs of vertices/clusters if there exists edges in *G*_ESG_ connecting vertices belonging to these two clusters respectively. The size of the vertex *i* corresponds to the entry of the count vector 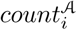. The entropy of the vector *count*𝒜 is also calculated to quantify how equally (or unequally) the algorithm 𝒜 samples graphs belonging to each cluster: 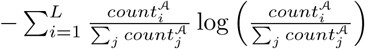. Larger entropy values indicate that the algorithm A samples each cluster more equally.

## 4 DISCUSSION

Over the last two decades, network motif discovery algorithms have been proposed that use several different underlying background graph sampling strategies. By all agreement in the literature, a uniform and independent background graph sampling method is fundamental for accurate network motif discovery, but evaluation of this condition on networks beyond tens of nodes was simply not possible because there was no feasible principle by which to perform such an evaluation. A number of works in the field of mathematical algorithms proposed methods that would provably sample uniformly within a defined error bound for nearly regular graphs ([37, 15]). However, most biological networks of interest for network motif finding studies, particularly gene regulatory networks, contain at least several hundred nodes and one or more “hubs” (for example, a transcription factor that is a master regulator). Thus, on the networks of biological interest, methods that used these mathematical approximation algorithms either did not apply due to regularity requirements on the network’s degree sequences, or the necessary computing time to achieve a result within a reasonable error bound rendered the method inapplicable. Direct uniformity tests were performed in the study of some algorithms by empirically enumerating all of the graphs in a small sample space, and this was helpful in the sense that it led to a general understanding in the field that graphs of uneven degree distribution posed significant problems for most algorithms. However, this left an extremely high level of uncertainty as to how these algorithms would perform in the case of realistic biological networks. As a result, despite the surge in popularity of network motif finding with the exciting findings reported in ([2, 4]), reported laboratory validations of predicted network motif instances are subsequently rare to nonexistent in multicellular organisms. With the IndeCut method, we hope to change this state of affairs by making it possible to evaluate the performance of any network motif finding algorithm on any network of interest, including biologically realistic networks.

In our study, we first demonstrate how IndeCut can be used by examining several major graph topologies and commonly used algorithms. By examining the performance of four published algorithms (FANMOD, DIA-MCIS, WaRSwap, and CoMoFinder) on both small example networks and large published biological networks from the literature (including E.coli and human regulatory networks), we conclude that different graph topologies can produce vast performance differences when using the same algorithm. We find that there are also large performance differences between algorithms on each graph topology. We then use IndeCut to compute how the performance of a sampling algorithm improves as the number of graphs in a sample increases. We examine where a performance plateau occurs for each graph and algorithm; we find that each algorithm does in fact have such a plateau, even if overall performance is not desirable. Alarmingly however, this plateau frequently occurs at a number of iterations far exceeding the recommended number of samples in program user manuals and/or defined by default software settings. Finally, we demonstrated that in cases of graphs with uneven degree distributions that are characteristic of biological networks, network motif discovery algorithms based on the graph randomization strategy known as edge-switching (the default strategy for the most commonly used algorithms) are vulnerable to highly non-uniform and/or non-independent sampling. Thus, this strategy is prone to spurious results on such networks. We use the concept of an edge-switching graph to show why this is the case. In essence, edge-switching algorithms produce a sampling bias by spending too much time sampling graphs that can be reached from the starting graph via a small number of edge-switches. Overall, IndeCut’s results show that some graph topologies are in fact highly troublesome to some algorithms (such as hub-containing graphs for edge-switch based algorithms), however, sampling performance cannot be anticipated universally for any algorithm or graph. For example, the WaRSwap algorithm handled hub-containing graphs with better performance as compared to other algorithms, while all algorithms except CoMoFinder had comparable performance on the near-regular graphs examined. CoMoFinder’s performance on near-regular graphs was closer to other algorithms on near-regular graphs than other graph types despite its overall performance difficulties. However in the case of “hybrid” graph topologies, algorithm performance varied significantly for each graph. In most cases, we observed that DIA-MCIS and WaRSwap both maintained relatively strong performance in sampling graphs with each example topology type; a similar trend in performance can be seen while evaluating the sampling performances on real biological networks. Nonetheless, algorithm performance variation across network types can be substantial. These results highlight IndeCut’s necessity in order to select an appropriate sampling algorithm for a biological network of interest.

The advent of IndeCut has significant implications for network motif finding studies that have been published in the literature over the last decade. These studies use the FANMOD program almost exclusively due to its early software availability and superior run times as compared with other programs available in the 2000’s. Studies performed using an edge-switching technique on small, relatively regular networks are likely justified. However, studies on enormous transcription factor-containing networks are very likely to contain spurious results in the sense that the outcomes obtained are not those that the user intended to measure. Since laboratory validation attempts on specific instances of network motifs are rarely if ever reported in these large network studies, the primary consequence is likely limited to spurious observations or conclusions that are only of general theoretical interest. By providing the community with an avenue to informed network motif discovery algorithm choice in the future, we hope that IndeCut will re-ignite interest in laboratory validation of the fascinating hypotheses that result from network motif discovery outcomes.

## 5 CONCLUSION

A fundamental component in the network motif discovery process is correct sampling of background networks. This process requires uniform and independent sampling from the space of all possible graphs given the degree sequence of the original network. Current network motif discovery algorithms use different heuristics in order to “shuffle” the original network, aiming to generate series of randomized networks which still preserve the original degree sequence. Without a sound mathematical approach for evaluating whether these “shuffled” collections of background graphs are generating uniform and independent samples, it was not possible to assess the accuracy of any network motif discovery outcome on a graph larger than tens of nodes. IndeCut provides such an approach, along with an implementation that is feasible for assessing algorithm performance on graphs containing tens of thousands of nodes and up to approximately one hundred thousand edges. Results from the application of IndeCut indicate that the most commonly used network motif discovery algorithm for background network randomization is likely to produce spurious outcomes on large realistic biological networks, and it is critically important to asses a network motif discovery algorithm’s accuracy on any network of interest if statistically justifiable conclusions are to be drawn from output. IndeCut provides an indispensable method for performing this assessment, particularly when costly laboratory validation of network motif discovery outcomes is under consideration.

## 6 ACKNOWLEDGEMENTS

We thank Jim Haglund of the University of Pennsylvania for early discussion of this effort with M.M., and pointing us to the recent work of Alexandar Barvinok of the University of Michigan in the field of enumerative combinatorics. We thank Alexander Barvinok for discussion of this work with D.K. M.M. and M.A. were supported by the NIH grant GM097188 to M.M.; and by startup funds from Oregon State University.

## 7 AVAILABILITY

IndeCut is an open source software and is available for download at https://github.com/megrawlab/IndeCut

## 8 Supplementary Data

### 8.1 Method S1: Description of examined network motif discovery algorithms

In order to compare the performance of existing network motif discovery algorithms using IndeCut, four different network motif finding algorithms were selected: FANMOD (Fast Network Motif Detection) [30], DIA-MCIS (Diaconis Monte Carlo Importance Sampling) [31], WaRSwap (Weighted and Reverse Swap sampling) [3], and CoMoFinder [32].

FANMOD is a well-known implementation of the edge switching randomization algorithm. The edge-switching method randomly chooses two directed edges (*x, y*), (*u, v*) from input graph *G* and switches their endpoints only if *G* doesn’t already contain either of these new edges (*x, v*), (*u, y*). It repeats this procedure for defined number of attempts and reports a random graph *G*′. An implemetation of FANMOD was downloaded from [30]. CoMoFinder implements a restricted version of the edge-switching method to detect only K-node motifs containing all node types such as TF, miRNA, and Gene, on given TF-miRNA-Gene regulatory networks. It breaks down the original network into seven different layers (miRNA → TF, TF → gene, miRNA TF, TF TF, TF → miRNA, TF → TF, TF → gene). Within each layer it chooses two edges (*x, y*), (*u, v*) and switches the endpoints if two conditions satisfied: 1) Neither of the new edge-pairs (*x, v*), (*u, y*) exist in the input graph *G*, and 2) An edge-switch between (*x, y*), (*u, v*) is allowed to happen only once, as revisiting a previously performed switch is not allowed (i.e. switching back from a graph containing (*x, v*) and (*u, y*) to a graph containing (*x, y*) and (*u, v*) is not allowed). CoMoFinder repeats the above-described procedure until either it reaches a stage such that no edge-pair is available to switch, or it has completed a pre-defined maximum number of edge-switching attempts. The original CoMoFinder program [32] was downloaded and modified to print randomized background graphs into files for our analysis.

DIA-MCIS is an efficient implementation of an importance sampling algorithm [19] to generate random graphs (self-loops included) from fixed in/out-degree sequences. DIA-MCIS converts an input graph *G* into a zero-one adjacency matrix *M*_*m*n*_ with *m* rows and *n* columns where *m*_*ij*_ is 1 if node *j* has a directed link to node *i*. It then sequentially fills the columns by a weighted-sampling scheme. It starts with first column which represents the first source node with out-degree of deg0, and assigns deg0 1s randomly to *m* cells (each cell represents a target node). In this process, nodes with higher in-degrees have more chance of selection by source nodes with higher out-degrees. The algorithm updates the row/column sums as proceeds to the next column.

WaRSwap produces randomized background graphs from an input graph by breaking it into layers representing five possible interaction types: TF→TF, TF→miRNA, TF→gene, miRNA→TF, and miRNA→gene. WaRSwap treats each layer as a bipartite graph *G* and operates as follows to generate a randomized graph *G*′. It first sorts the source nodes in descending order of out-degree, and for each source node *S*_*i*_ it computes the sampling weights for each target node *T*_*j*_ using a weighting formula [3]. The weighting formula corrects the tendency of source nodes with large out-degrees to target nodes with larger in-degrees. WaRSwap places an edge between *S*_*i*_ and *T*_*j*_ if possible, otherwise it enters a specific back-swapping procedure to identify a new target node. We downloaded a Java implementation of WaRSwap from http://megraw.cgrb.oregonstate.edu/software/WaRSwapSoftwareApplication/ and R implementation from http://megraw.cgrb.oregonstate.edu/software/WaRSwap. The WaRSwapApp makes an automated selection of the WaRSwap weighting parameter for the user based on the in/out-degree sequences of the input graph. We modified the R implementation of WaRSwap to include this automated weighting parameter selection.

### 8.2 Supplementary Figures

**Figure S1:**
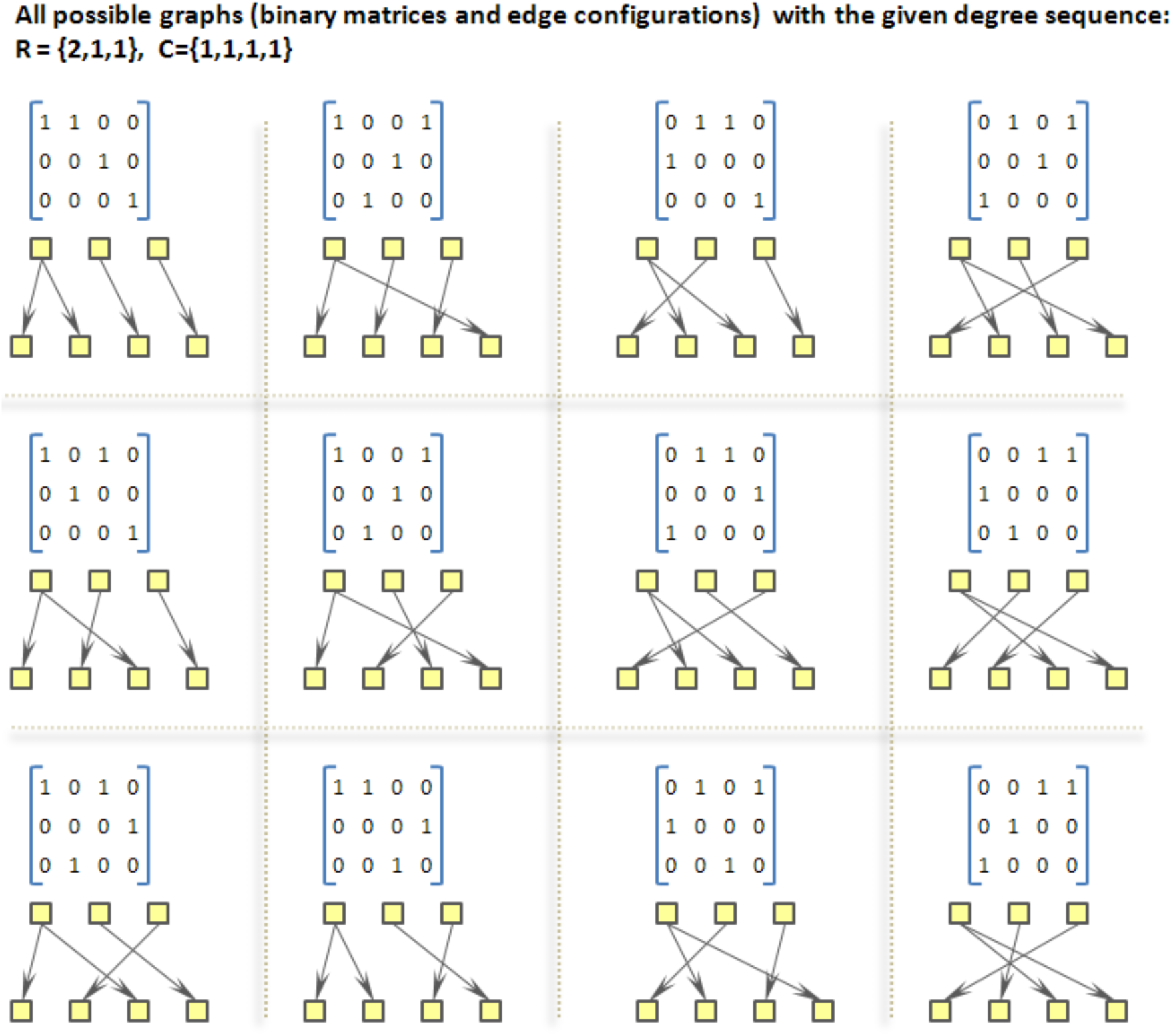
Sample space of an example degree sequence.The sample space of this degree sequence contains 12 different graphs.

**Figure S2:**
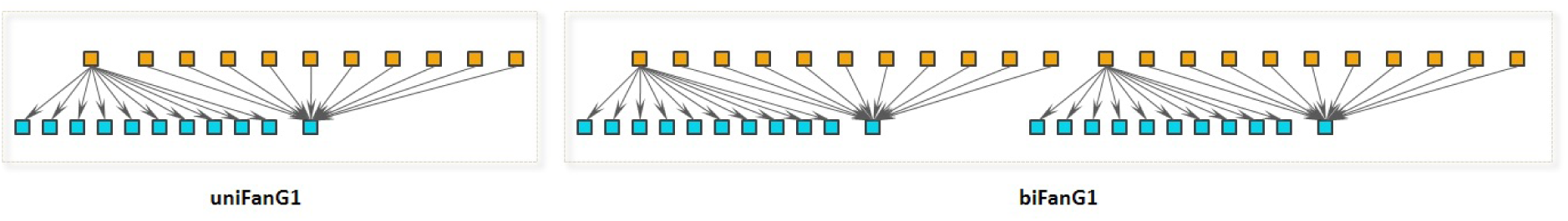
Constructing multiFan graphs starting from uniFanG1. A biFan graph is created by attaching two uniFanG1 graphs.

**Figure S3:**
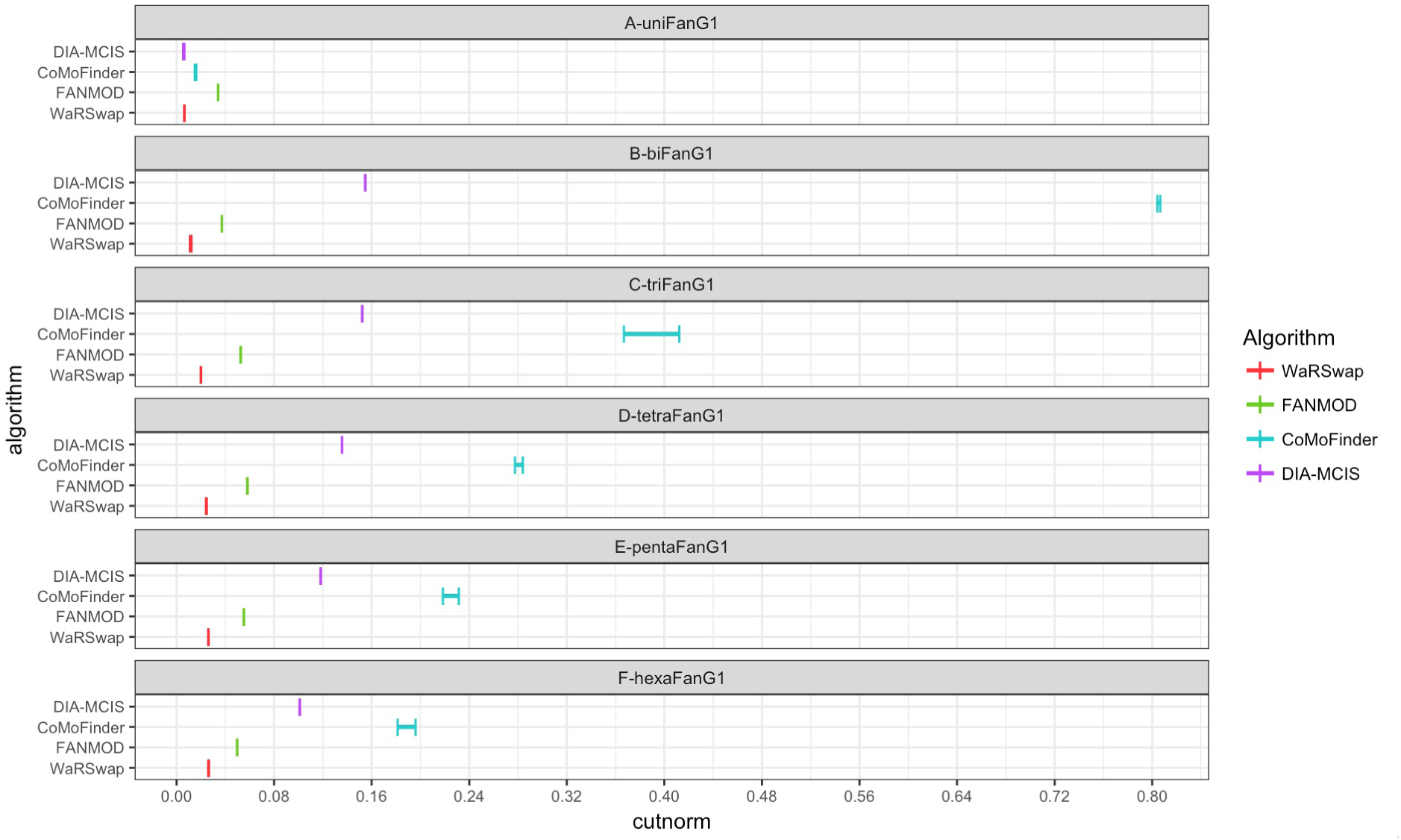
Graph sampling performance evaluation on small uneven graphs using IndeCut. This figure shows the cut norm estimates for all four examined algorithms: WaRSwap, CoMoFinder, DIA-MCIS, and FANMOD. For each graph and algorithm, 5000 graphs were generated. The cut norm estimates for each algorithm were computed using IndeCut. A smaller cut norm interval that is closer to zero represents more uniform and independent sampling.

**Figure S4:**
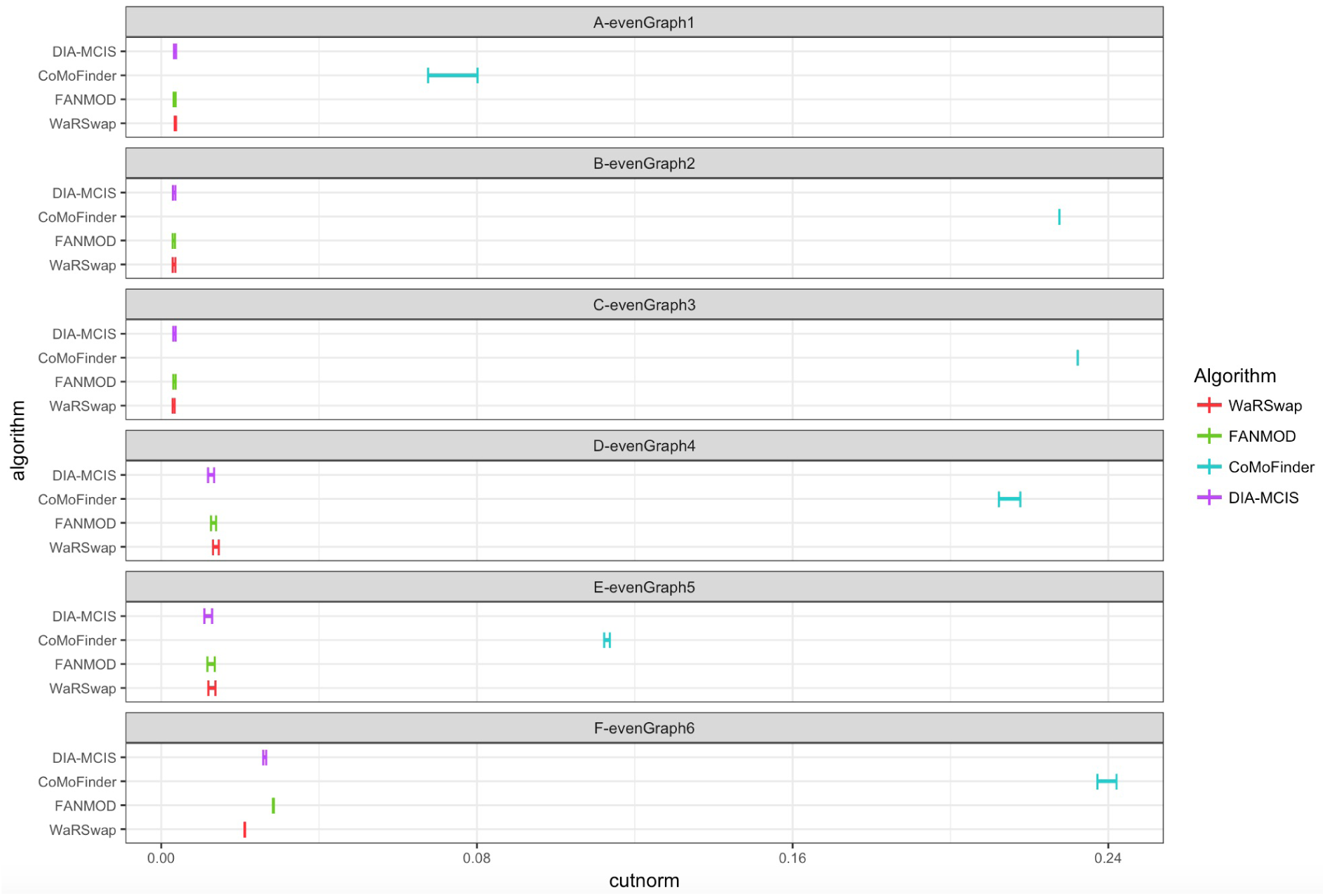
Graph sampling performance evaluation on small even graphs using IndeCut. This figure shows the cut norm estimates for all four examined algorithms: WaRSwap, CoMoFinder, DIA-MCIS, and FANMOD. For each graph and algorithm, 5000 graphs were generated. The cut norm estimates for each algorithm were computed using IndeCut. A smaller cut norm interval that is closer to zero represents more uniform and independent sampling.

**Figure S5:**
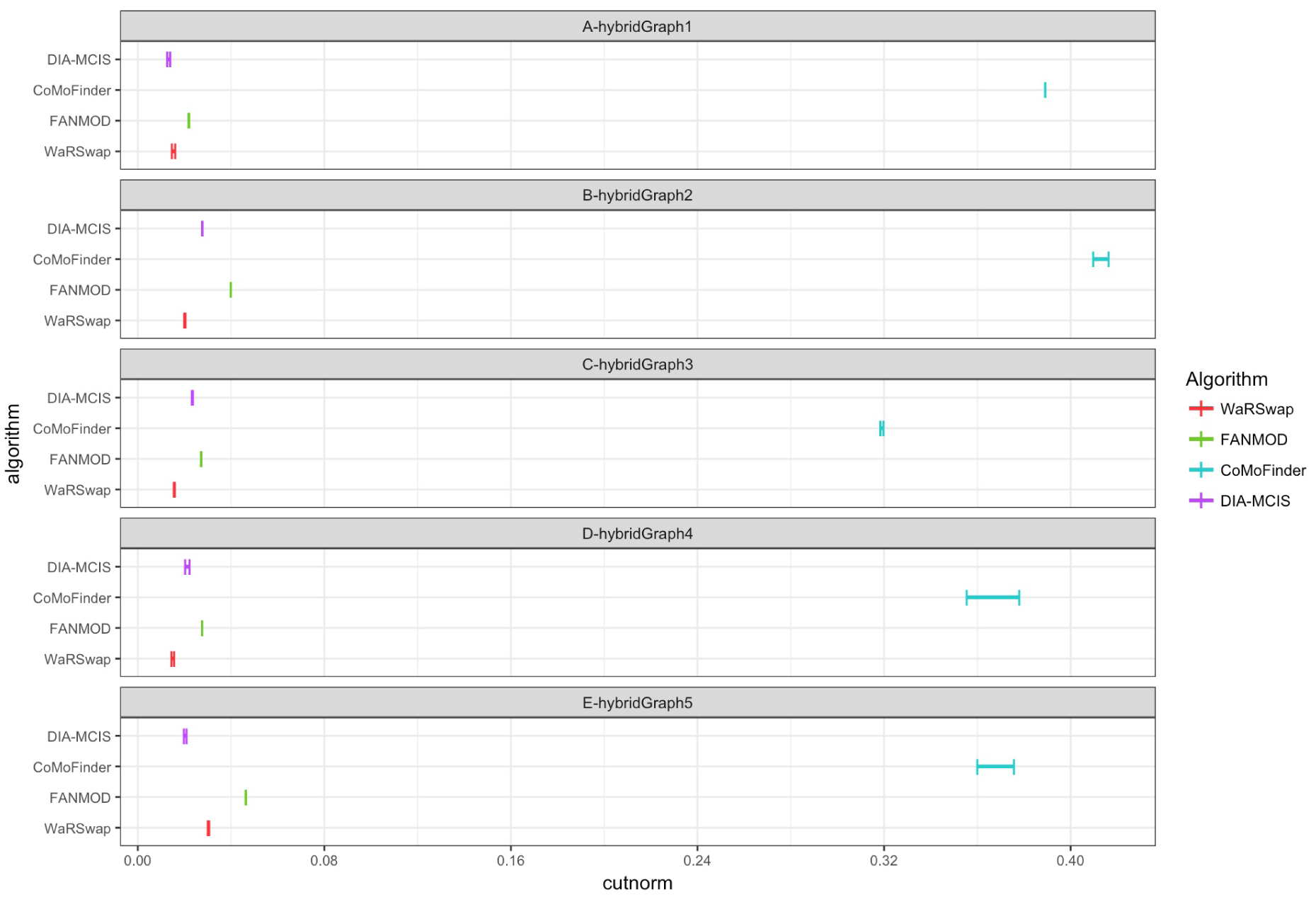
Graph sampling performance evaluation on small hybrid graphs using IndeCut. This figure shows the cut norm estimates for all four examined algorithms: WaRSwap, CoMoFinder, DIA-MCIS, and FANMOD. For each graph and algorithm, 5000 graphs were generated. The cut norm estimates for each algorithm were computed using IndeCut. A smaller cut norm interval that is closer to zero represents more uniform and independent sampling.

**Figure S6:**
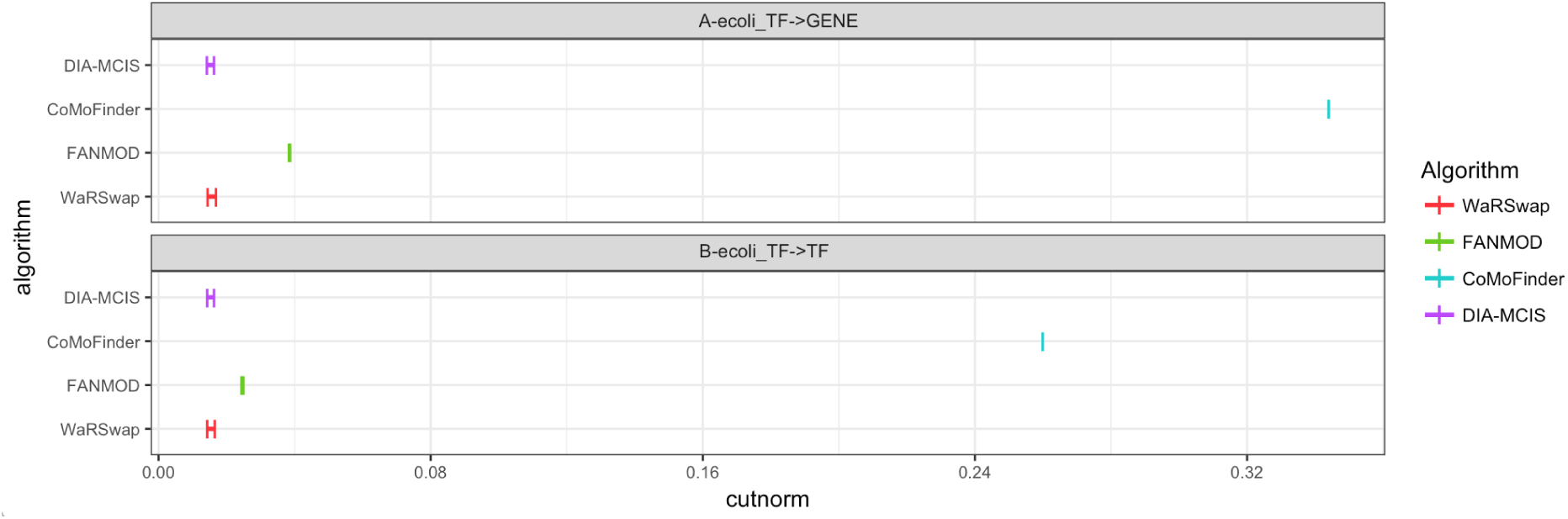
Graph sampling performance evaluation on Ecoli network using IndeCut. This figure shows the cut norm estimates for all four examined algorithms: WaRSwap, CoMoFinder, DIA-MCIS, and FANMOD. For each graph and algorithm, 5000 graphs were generated. The cut norm estimates for each algorithm were computed using IndeCut. A smaller cut norm interval that is closer to zero represents more uniform and independent sampling.

**Figure S7:**
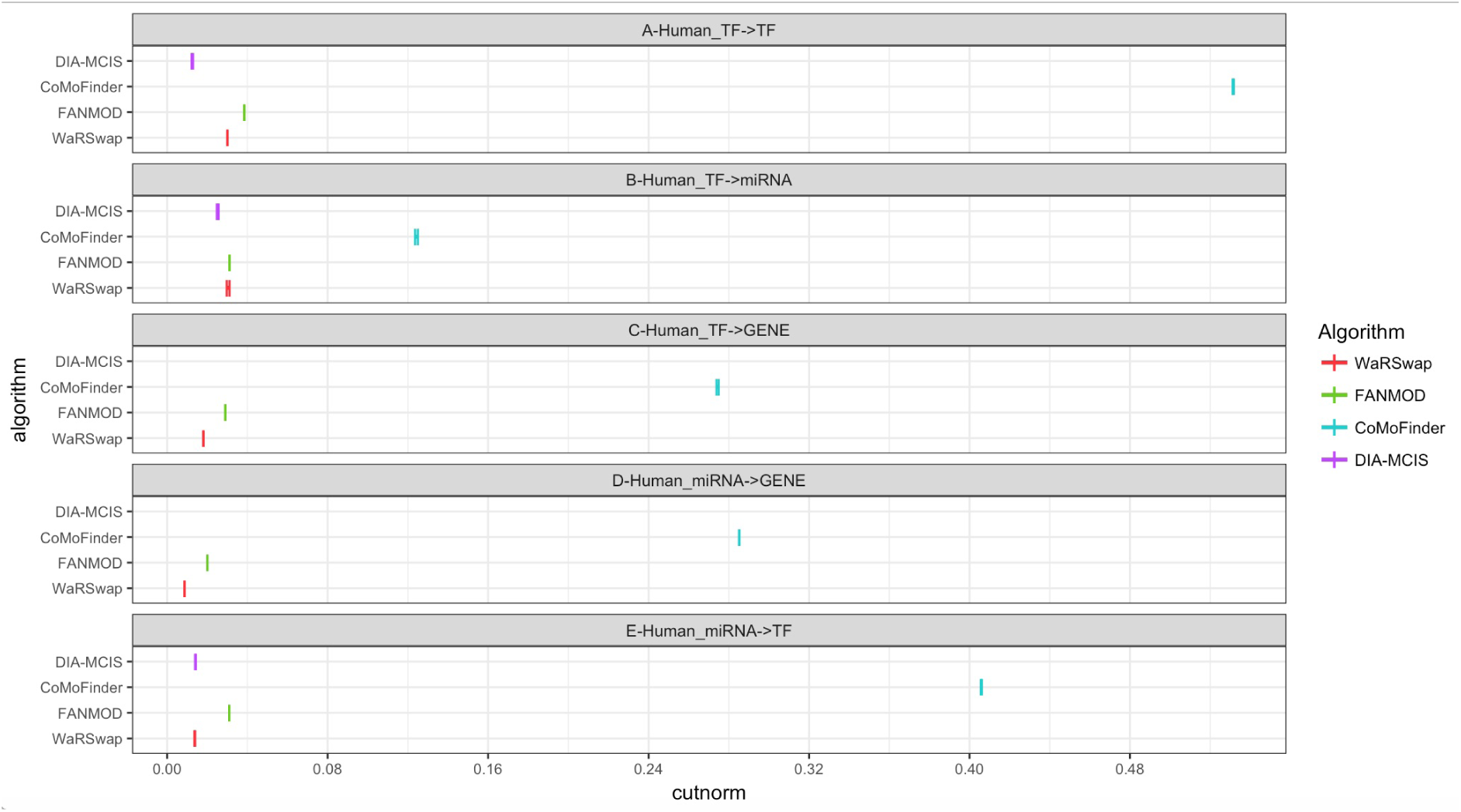
Graph sampling performance evaluation on Human regulatory network using IndeCut. This figure shows the cut norm estimates for all four examined algorithms: WaRSwap, CoMoFinder, DIA-MCIS, and FANMOD. The cut norm estimates for DIA-MCIS are absent from C and D because this algorithm is not able to perform on large graphs with more that 2,035 nodes.

**Figure S8:**
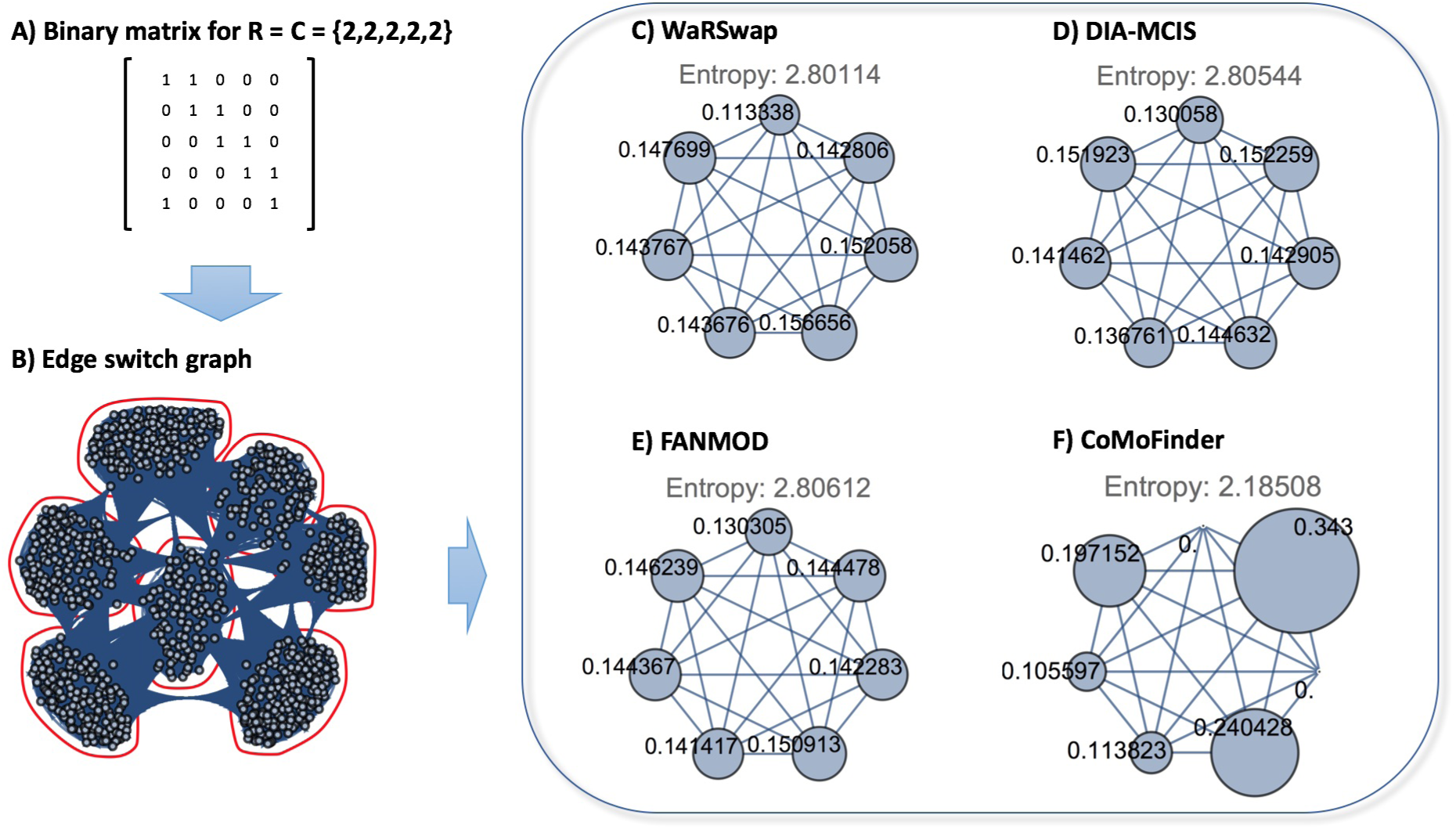
The ESG graph and cluster-time diagrams for an example even graph. A) The zero-one matrix representation of an even graph with degree sequence of R=C= {2,2,2,2,2}. B) The ESG graph corresponding to the graph in part A. Running the graph clustering algorithm on the ESG graph detects seven different clusters. C-F) The cluster-time diagrams for each examined algorithm were computed and visualized.

**Figure S9:**
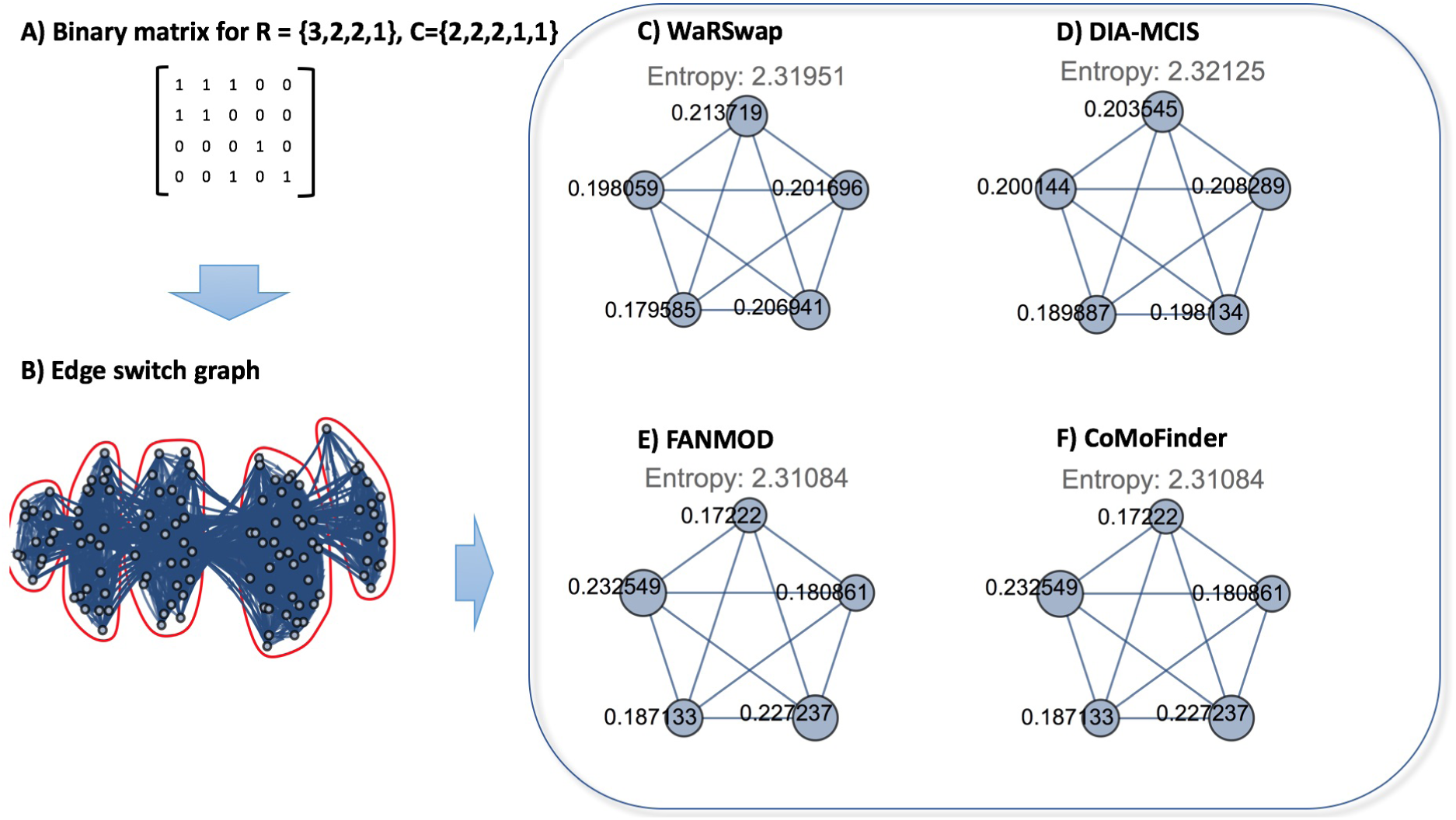
The ESG graph and cluster-time diagrams for an example hybrid graph. A) The zero-one matrix representation of an uneven graph with degree sequence of R= {3,2,2,1}, C= {2,2,2,1,1}. B) The ESG graph corresponding to the graph in part A. Running the graph clustering algorithm on the ESG graph detects five different clusters. C-F) The cluster-time diagrams for each examined algorithm were computed and visualized.

**Figure S10:**
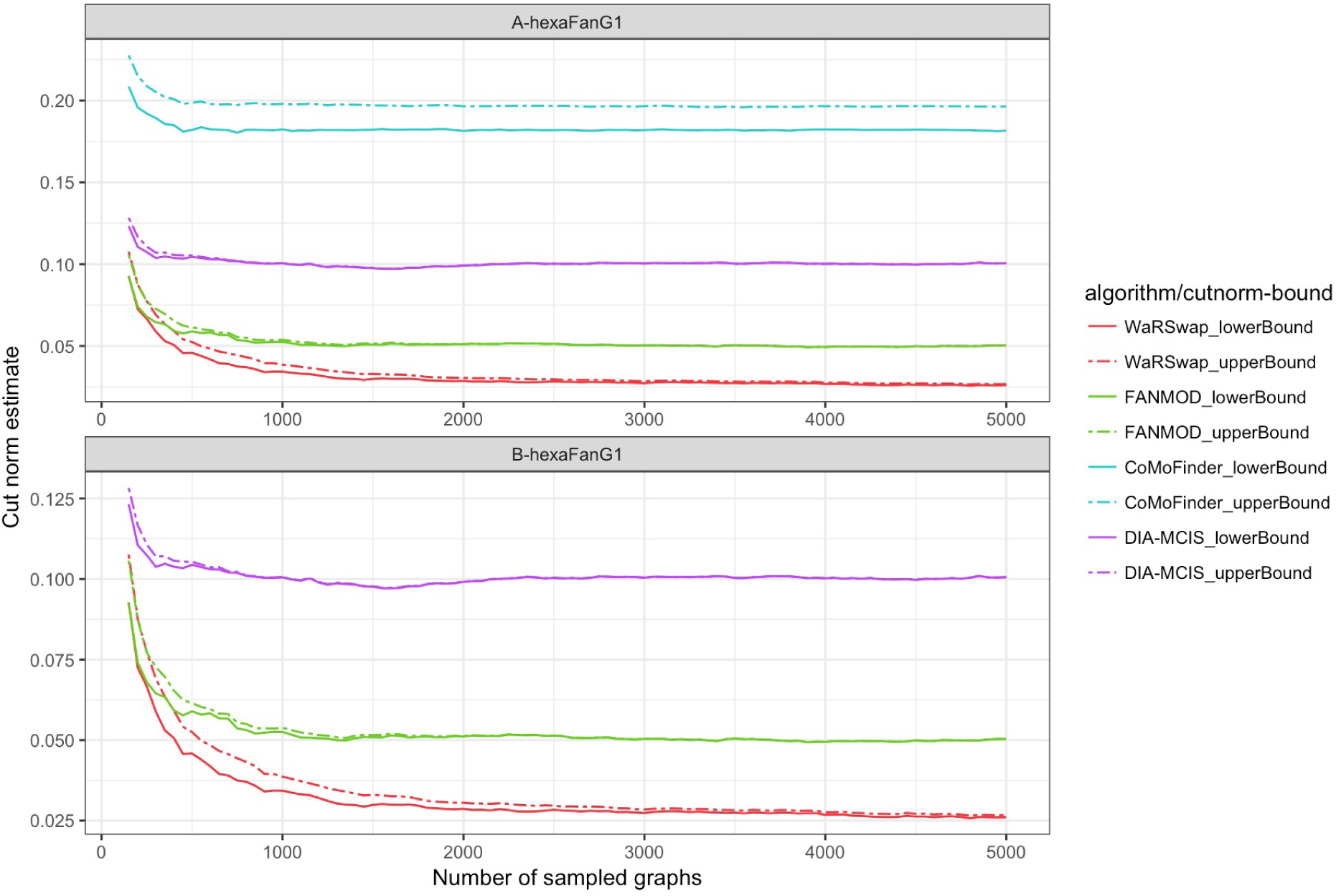
Sampling performance vs. the number of samples for graph hexaFanG1. All 5000 samples previously generated by each algorithm for hexaFanG1 were collected and subsampled into 25 sets (200, 400, 600,, 5000 samples in each set, respectively). IndeCut was used to compute the cut norm estimates (lower and upper bounds) for each set of samples and algorithms. Cut norm values closer to zero represent a more uniform/independent sampling. A) The relationship between the sampling performance and number of samples for all four examined algorithms is shown. B) The relationship between the sampling performance and number of samples for three algorithms WaRSwap, FANMOD, and DIA-MCIS is shown. The cut norm estimates for CoMoFinder were removed from this figure for ease of comparison (CoMoFinder has much larger cut norm estimates as compared to other three algorithms).

**Figure S11:**
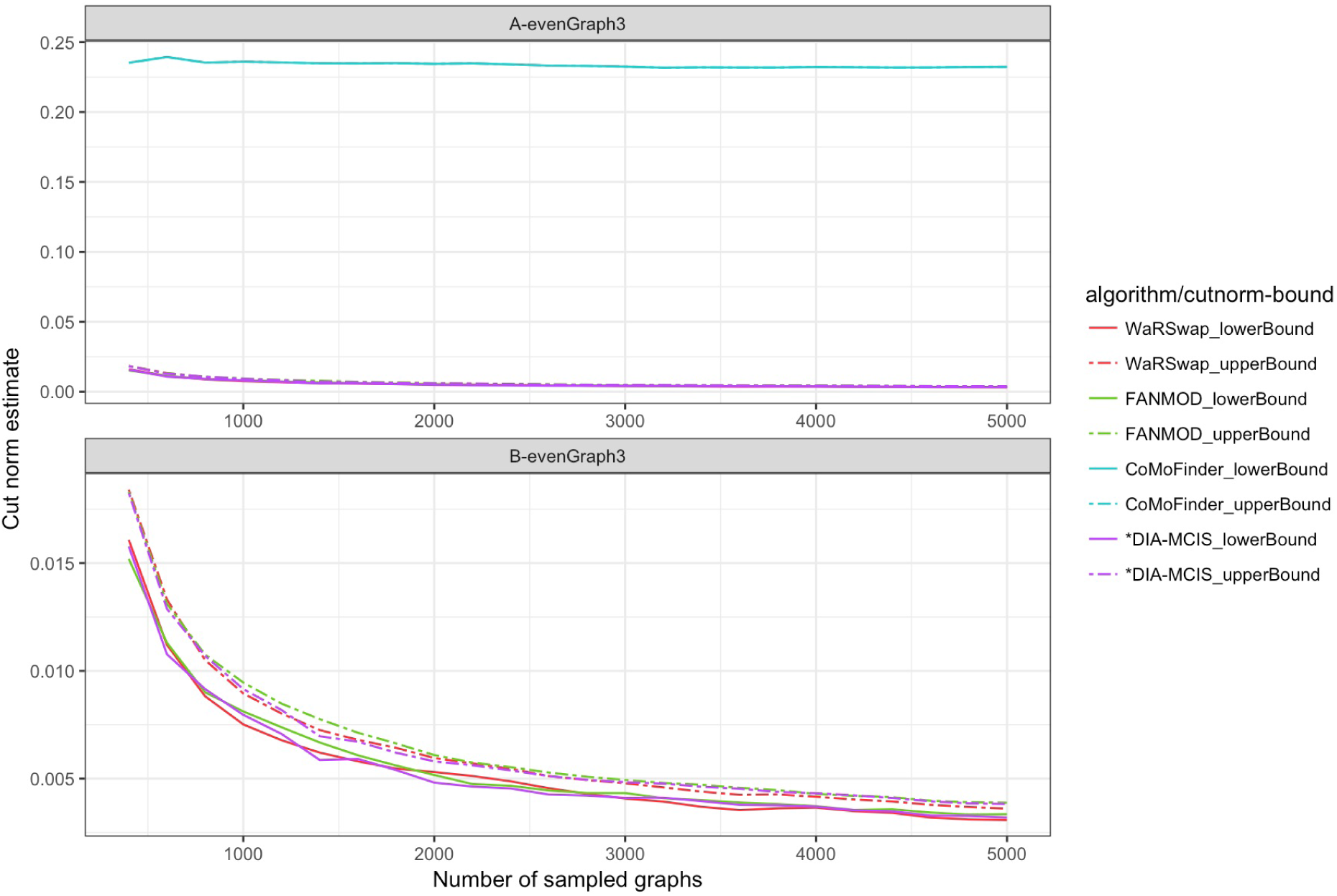
The cut norm estimates vs. the number of samples for graph evenGraph3. All 5000 samples previously generated by each algorithm for evenGraph3 network were collected and subsampled into 25 sets (200, 400, 600,, 5000 samples in each set, respectively). IndeCut was used to compute the cut norm estimates (lower and upper bounds) for each set of samples and algorithms. Cut norm values closer to zero represent a more uniform/independent sampling. A) The relationship between the sampling performance and number of samples for all four examined algorithms is shown. B) The relationship between the sampling performance and number of samples for three algorithms WaRSwap, FANMOD, and DIA-MCIS is shown. The cut norm estimates for CoMoFinder were removed from this figure for ease of comparison (CoMoFinder has much larger cut norm estimates as compared to other three algorithms).

**Figure S12:**
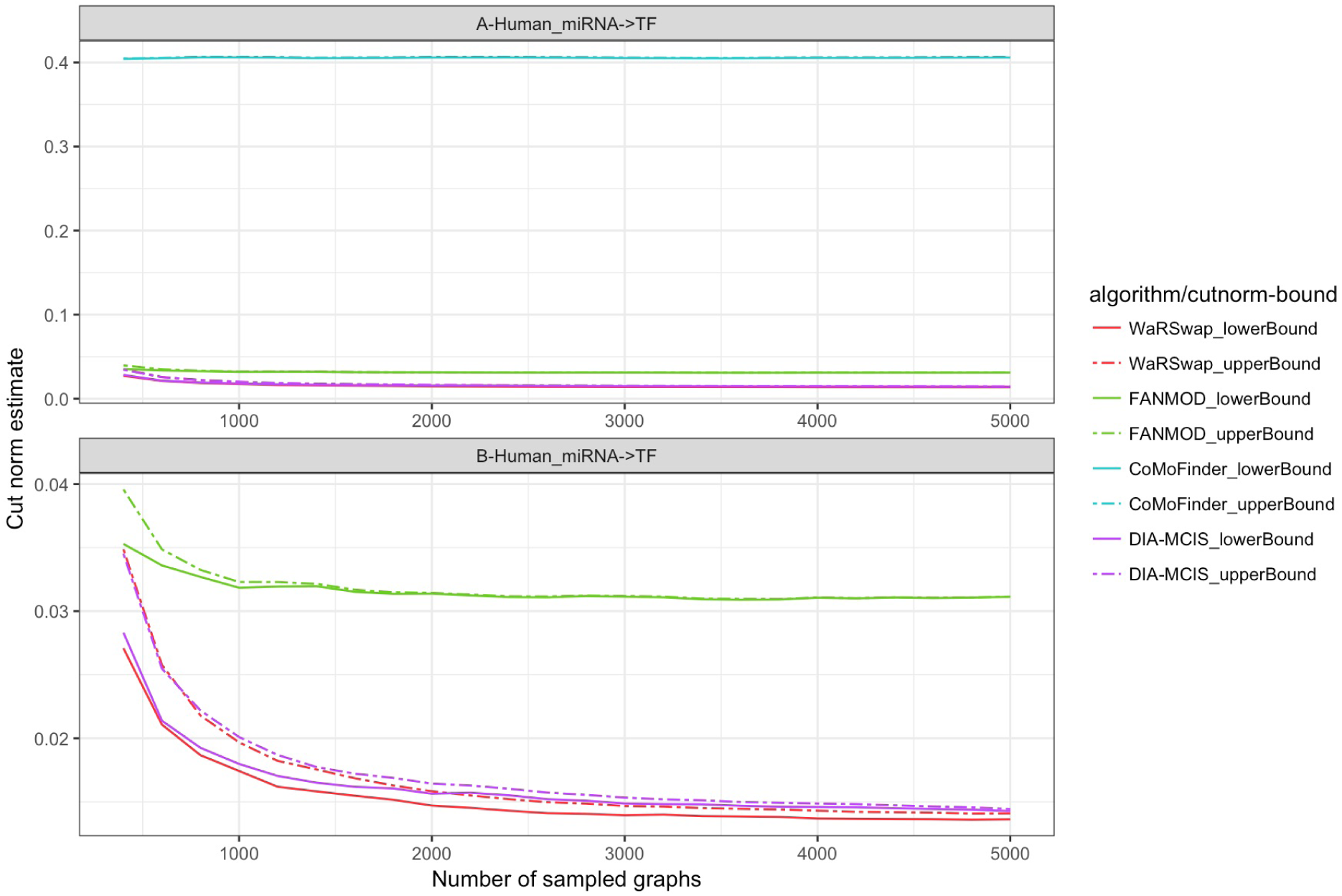
The cut norm estimates vs. the number of samples for Human miRNA→TF network. All 5000 samples previously generated by each algorithm for Human miRNA→TF network were collected and subsampled into 25 sets (200, 400, 600,, 5000 samples in each set, respectively). IndeCut was used to compute the cut norm estimates (lower and upper bounds) for each set of samples and algorithms. Cut norm values closer to zero represent a more uniform/independent sampling. A) The relationship between the sampling performance and number of samples for all four examined algorithms is shown. B) The relationship between the sampling performance and number of samples for three algorithms WaRSwap, FANMOD, and DIA-MCIS is shown. The cut norm estimates for CoMoFinder were removed from this figure for ease of comparison (CoMoFinder has much larger cut norm estimates as compared to other three algorithms).

**Table S1:** Table of cut norm estimates for all examined graphs. The cut norm estimates closer to zero represents more uniform and independent sampling.

| graphName | no_samples | Z_low<br>cutnorm | Z_up<br>cutnorm | WR_low<br>cutnorm | WR_up<br>cutnorm | FN_low<br>cutnorm | FN_up<br>cutnorm | comoF_low<br>cutnorm | comoF_up<br>cutnorm | diamcis_low<br>cutnorm | diamcis_up<br>cutnorm |
| --- | --- | --- | --- | --- | --- | --- | --- | --- | --- | --- | --- |
| uniFanG1 | 5000 | 5.000004 | 5.000004 | 0.0065 | 0.0066 | 0.0343 | 0.0343 | 0.015 | 0.0164 | 0.0057 | 0.0064 |
| biFanG1 | 5000 | 10.000006 | 10.000006 | 0.0111 | 0.0122 | 0.0372 | 0.0373 | 0.8042 | 0.8068 | 0.1549 | 0.1549 |
| triFanG1 | 5000 | 14.999994 | 14.999994 | 0.0198 | 0.0201 | 0.0527 | 0.0528 | 0.367 | 0.4123 | 0.1525 | 0.1525 |
| tetraFanG1 | 5000 | 20.000003 | 20.000003 | 0.0241 | 0.0248 | 0.0582 | 0.0583 | 0.2778 | 0.2842 | 0.1358 | 0.1358 |
| pentaFanG1 | 5000 | 24.999999 | 24.999999 | 0.0259 | 0.0264 | 0.0552 | 0.0552 | 0.2185 | 0.2315 | 0.1182 | 0.1182 |
| F-hexaFanG1 | 5000 | 29.999998 | 29.999998 | 0.0261 | 0.0266 | 0.0498 | 0.0499 | 0.1814 | 0.1962 | 0.1011 | 0.1011 |
| evenGraph1 | 5000 | 200.000013 | 200.000013 | 0.0034 | 0.0038 | 0.0032 | 0.0037 | 0.0677 | 0.0802 | 0.0033 | 0.0038 |
| evenGraph2 | 5000 | 264.499966 | 264.5 | 0.0029 | 0.0036 | 0.0029 | 0.0035 | 0.2276 | 0.2276 | 0.003 | 0.0036 |
| evenGraph3 | 5000 | 242.000044 | 242.000044 | 0.0029 | 0.0034 | 0.0031 | 0.0037 | 0.2323 | 0.2323 | 0.0031 | 0.0037 |
| evenGraph4 | 5000 | 5.5 | 5.5 | 0.0132 | 0.0146 | 0.0127 | 0.0139 | 0.2123 | 0.2177 | 0.0119 | 0.0134 |
| evenGraph5 | 5000 | 21.499986 | 21.499987 | 0.012 | 0.0138 | 0.0117 | 0.0136 | 0.1123 | 0.1137 | 0.011 | 0.0129 |
| evenGraph6 | 5000 | 7.999996 | 7.999996 | 0.0211 | 0.0213 | 0.0283 | 0.0286 | 0.2373 | 0.2421 | 0.0259 | 0.0266 |
| hybridGraph1 | 5000 | 9.499999 | 9.499999 | 0.0147 | 0.0161 | 0.0218 | 0.0221 | 0.3892 | 0.3892 | 0.0126 | 0.0139 |
| hybridGraph2 | 5000 | 96.750016 | 96.750016 | 0.0199 | 0.0205 | 0.0399 | 0.0399 | 0.4097 | 0.4163 | 0.0277 | 0.0278 |
| hybridGraph3 | 5000 | 111.000018 | 111.000018 | 0.0155 | 0.016 | 0.0272 | 0.0272 | 0.3184 | 0.3197 | 0.0232 | 0.0237 |
| hybridGraph4 | 5000 | 42.499996 | 42.499996 | 0.0145 | 0.0157 | 0.0276 | 0.0276 | 0.3555 | 0.378 | 0.0204 | 0.0222 |
| hybridGraph5 | 5000 | 20.000008 | 20.000008 | 0.0301 | 0.0306 | 0.0463 | 0.0464 | 0.36 | 0.3757 | 0.0198 | 0.0209 |
| ecoli_TF-GENE | 5000 | 97.499958 | 97.498955 | 0.0145 | 0.0169 | 0.0384 | 0.0388 | 0.344 | 0.3441 | 0.0142 | 0.0164 |
| ecoli_TF-TF | 5000 | 32.249996 | 32.250064 | 0.0144 | 0.0166 | 0.0244 | 0.025 | 0.26 | 0.26 | 0.0143 | 0.0163 |
| Human_TF→TF | 5000 | 161.000002 | 161.000002 | 0.0301 | 0.0301 | 0.0385 | 0.0385 | 0.5313 | 0.5318 | 0.0123 | 0.0129 |
| Human_TF→miRNA | 5000 | 309.250145 | 309.250145 | 0.0297 | 0.0312 | 0.0312 | 0.0312 | 0.1236 | 0.125 | 0.0248 | 0.0258 |
| Human_TF→GENE | 5000 | 6437.000388 | 6437.000388 | 0.0181 | 0.0183 | 0.029 | 0.029 | 0.274 | 0.275 |  |  |
| Human_miRNA→GENE | 5000 | 28855.25551 | 28855.25551 | 0.0087 | 0.0088 | 0.0202 | 0.0202 | 0.2852 | 0.2853 |  |  |
| Human_miRNA→TF | 5000 | 648.50001 | 648.50001 | 0.0136 | 0.014 | 0.031 | 0.031 | 0.4057 | 0.4062 | 0.0141 | 0.0143 |

**Table S2:** Runtime of IndeCut on all examined graphs. IndeCut evaluates graphs on the order of several thousand nodes and tens of thousands of edges within a few minutes to a few days using standard hardware. This table provides IndeCut’s observed run time on each graph and algorithm. The miRNA→Gene layer in the human network allows us to provide run time given an extreme example with approximately 100,000 edges. To put these run times into perspective, network motif tools typically take several days simply to provide an output for graphs of this size, using a small number of iterations that does not guarantee meaningfully accurate performance (we discuss the number of iterations necessary for optimal performance for each sampling method in the next section). Using a commercial optimization package such as Guorbi or Mosek (in contrast to the open-source package CSDP that we use here) will result in speed improvements to IndeCut. Thus, considering time costs of running network motif finding algorithms themselves as well as the enormous potential laboratory costs of attempting to validate inaccurate results, IndeCut presents a very practical method for making an informed network motif discovery algorithm choice on biological networks of study.

| Graph | Number of nodes | Number of edges | IndeCut run-time |
| --- | --- | --- | --- |
| uniFanG1 | 11 | 20 | 10 s |
| biFanG1 | 22 | 40 | 10 s |
| triFanG1 | 33 | 60 | 15 s |
| tetraFanG1 | 44 | 80 | 20 s |
| pentaFanG1 | 55 | 100 | 35 s |
| hexaFanG1 | 66 | 120 | 48 s |
| regularSmallG1 | 23 | 22 | 9 s |
| regularSmallG2 | 62 | 86 | 21 s |
| regularSmallG3 | 31 | 32 | 10 s |
| regularG1 | 80 | 800 | 27 s |
| regularG2 | 92 | 1058 | 30 s |
| regularG3 | 88 | 968 | 28 s |
| Human_TF→TF | 174 | 644 | 52 s |
| Human_TF→miR | 332 | 1237 | 2 min |
| Human_miR→TF | 606 | 2594 | 5 min |
| Human_TF→GENE | 9055 | 25748 | 4 days |
| Human_miR→GENE | 12185 | 115421 | 14 days |
| Ecoli_TF→TF | 140 | 129 | 2 min |
| Ecoli_TF→GENE | 365 | 390 | 4 min |

## Reference

[1] Gaudinier, A. and Brady, S. M. (2016) Mapping Transcriptional Networks in Plants: Data-Driven Discovery of Novel Biological Mechanisms. Annual review of plant biology, 67, 575–594.

[2] Milo, R., Shen-Orr, S., Itzkovitz, S., Kashtan, N., Chklovskii, D., and Alon, U. (2002) Network motifs: simple building blocks of complex networks. Science, 298(5594), 824–827.

[3] Megraw, M., Mukherjee, S., and Ohler, U. (2013) Sustained-input switches for transcription factors and microRNAs are central building blocks of eukaryotic gene circuits. Genome biology, 14(8), 1.

[4] Alon, U. (2007) Network motifs: theory and experimental approaches. Nature Reviews Genetics, 8(6), 450–461.

[5] Wong, E., Baur, B., Quader, S., and Huang, C.-H. (2011) Biological network motif detection: principles and practice. Briefings in bioinformatics, p. bbr033.

[6] Mangan, S. and Alon, U. (2003) Structure and function of the feed-forward loop network motif. Proceedings of the National Academy of Sciences, 100(21), 11980–11985.

[7] Shen-Orr, S. S., Milo, R., Mangan, S., and Alon, U. (2002) Network motifs in the transcriptional regulation network of Escherichia coli. Nature genetics, 31(1), 64–68.

[8] Barabasi, A.-L. and Oltvai, Z. N. (2004) Network biology: understanding the cell’s functional organization. Nature reviews genetics, 5(2), 101–113.

[9] Wang, P., Lü, J., Yu, X., and Liu, Z. (2015) Duplication and divergence effect on network motifs in undirected bio-molecular networks. IEEE transactions on biomedical circuits and systems, 9(3), 312–320.

[10] Ribeiro, P., Silva, F., and Kaiser, M. (2009) Strategies for network motifs discovery. In e-Science, 2009. e-Science’09. Fifth IEEE International Conference IEEE pp. 80–87.

[11] Tran, N. T. L., DeLuccia, L., McDonald, A. F., and Huang, C.-H. (2015) Cross-disciplinary detection and analysis of network motifs. Bioinformatics and Biology insights, 9, 49.

[12] Milo, R., Kashtan, N., Itzkovitz, S., Newman, M. E., and Alon, U. (2003) On the uniform generation of random graphs with prescribed degree sequences. arXiv preprint cond-mat/0312028,.

[13] Barabási, A.-L. and Albert, R. (1999) Emergence of scaling in random networks. science, 286(5439), 509–512.

[14] Fosdick, B. K., Larremore, D. B., Nishimura, J., and Ugander, J. (2016) Configuring Random Graph Models with Fixed Degree Sequences. arXiv preprint arXiv:1608.00607,.

[15] Bezáková, I., Bhatnagar, N., and Vigoda, E. (2007) Sampling binary contingency tables with a greedy start. Random Structures & Algorithms, 30(1-2), 168–205.

[16] Chatterjee, S., Diaconis, P., and Sly, A. (2011) Random graphs with a given degree sequence. The Annals of Applied Probability, pp. 1400–1435.

[17] Erdös, P. L., Miklós, I., and Soukup, L. (2010) Towards random uniform sampling of bipartite graphs with given degree sequence. arXiv preprint arXiv:1004.2612,.

[18] Itzkovitz, S., Milo, R., Kashtan, N., Ziv, G., and Alon, U. (2003) Subgraphs in random networks. Physical review E, 68(2), 026127.

[19] Chen, Y., Diaconis, P., Holmes, S. P., and Liu, J. S. (2005) Sequential Monte Carlo methods for statistical analysis of tables. Journal of the American Statistical Association, 100(469), 109–120.

[20] King, O. D. (2004) Comment on Subgraphs in random networks. Physical Review E, 70(5), 058101.

[21] Kim, W., Diko, M., and Rawson, K. (2013) Network motif detection: Algorithms, parallel and cloud computing, and related tools. Tsinghua Science and Technology, 18(5), 469–489.

[22] Thomas, S. and Bonchev, D. (2010) A survey of current software for network analysis in molecular biology. Human genomics, 4(5), 1.

[23] Grochow, J. A. and Kellis, M. (2007) Network motif discovery using subgraph enumeration and symmetry-breaking. In Annual International Conference on Research in Computational Molecular Biology Springer pp. 92–106.

[24] Blitzstein, J. and Diaconis, P. (2011) A sequential importance sampling algorithm for generating random graphs with prescribed degrees. Internet Mathematics, 6(4), 489–522.

[25] Greenhill, C. (2015) The switch Markov chain for sampling irregular graphs. In Proceedings of the Twenty-Sixth Annual ACM-SIAM Symposium on Discrete Algorithms SIAM pp. 1564–1572.

[26] Sorrells, T. R. and Johnson, A. D. (2015) Making sense of transcription networks. Cell, 161(4), 714–723.

[27] Winterbach, W., Van Mieghem, P., Reinders, M., Wang, H., and de Ridder, D. (2013) Topology of molecular interaction networks. BMC systems biology, 7(1), 1.

[28] Alon, N. and Naor, A. (2006) Approximating the cut-norm via Grothendieck’s inequality. SIAM Journal on Computing, 35(4), 787–803.

[29] Barvinok, A. (2010) On the number of matrices and a random matrix with prescribed row and column sums and 0–1 entries. Advances in Mathematics, 224(1), 316–339.

[30] Wernicke, S. and Rasche, F. (2006) FANMOD: a tool for fast network motif detection. Bioinformatics, 22(9), 1152–1153.

[31] Fusco, D., Bassetti, B., Jona, P., and Lagomarsino, M. C. (2007) DIA-MCIS: an importance sampling network randomizer for network motif discovery and other topological observables in transcription networks. Bioinformatics, 23(24), 3388–3390.

[32] Liang, C., Li, Y., Luo, J., and Zhang, Z. (2015) A novel motif-discovery algorithm to identify co-regulatory motifs in large transcription factor and microRNA co-regulatory networks in human. Bioinformatics, 31(14), 2348–2355.

[33] Roy, S., Ernst, J., Kharchenko, P. V., Kheradpour, P., Negre, N., Eaton, M. L., Landolin, J. M., Bristow, C. A., Ma, L., Lin, M. F., et al. (2010) Identification of functional elements and regulatory circuits by Drosophila modENCODE. Science, 330(6012), 1787–1797.

[34] Brandes, U., Delling, D., Gaertler, M., Gorke, R., Hoefer, M., Nikoloski, Z., and Wagner, D. (2008) On modularity clustering. IEEE Transactions on Knowledge and Data Engineering, 20(2), 172–188.

[35] Borchers, B. (1999) CSDP, AC library for semidefinite programming. Optimization methods and Software, 11(1-4), 613–623.

[36] Janson, S., Graphons, cut norm and distance, couplings and rearrangements. Technical report, Department of Mathematics, Uppsala University (2010).

[37] Bayati, M., Kim, J. H., and Saberi, A. (2010) A sequential algorithm for generating random graphs. Algorithmica, 58(4), 860–910.

